# Mechanical Signaling Drives Tunneling Nanotubes to Preserve Cytoskeleton Tension and Lamin Integrity Against α-Synuclein-Induced Senescence in Astroglia

**DOI:** 10.64898/2026.03.13.711517

**Authors:** Suchana Chatterjee, Ashok Ravula, Anirudh Sreenivas BK, Abinaya Raghavan, Niharika Shivanandaswamy, Sivaraman Padavattan, Sreenath Balakrishnan, Sangeeta Nath

## Abstract

Astroglia can counteract the harmful effects of α-synuclein (α-SYN) protofibrils and reverse premature cellular senescence by promoting tunneling nanotubes (TNTs). However, the mechanism behind this recovery is unknown. This study is the first to examine TNT–mediated mechanical stability in senescent astroglial recovery. We demonstrate that disruption of Lamin A/C in α-SYN-protofibrils-treated senescent cells reduces actin-cytoskeleton stress, as measured by nucleus flatness index and isometric scale factor from quantitative microscopy. ROCK (Rho-associated kinase) inhibition, which is crucial for reducing actin-cytoskeleton tension, promotes TNTs. Small molecules like Cytochalasin-D, Nocodazole, and Jasplakinolide, which inhibit TNTs by altering actin tension other than ROCK pathway, cannot reverse senescence. RNA-sequence heatmaps reveal changes in senescence-, integrin-, and ROCK-pathway genes; STRING links these to the Hippo pathway. Experimental results show that cytosolic YAP translocation, a key regulator of Hippo pathway, is vital for TNT formation and actin-based stability in U87-MG astrocytoma and primary astrocytes. Interestingly, TNTs form between two cells with different actin tensions: one exhibits low actin tension with Hippo signaling on, while the other has higher actin tension with Hippo signaling off. The most notable observation is the high abundance of YAP inside the TNTs, along with actin. The study shows that TNTs maintain mechanical stability through Lamin A/C integrity and actin tension in α-SYN-induced senescent astroglia, thereby protecting the cells, reversing senescence.

**Graphical Abstract:** 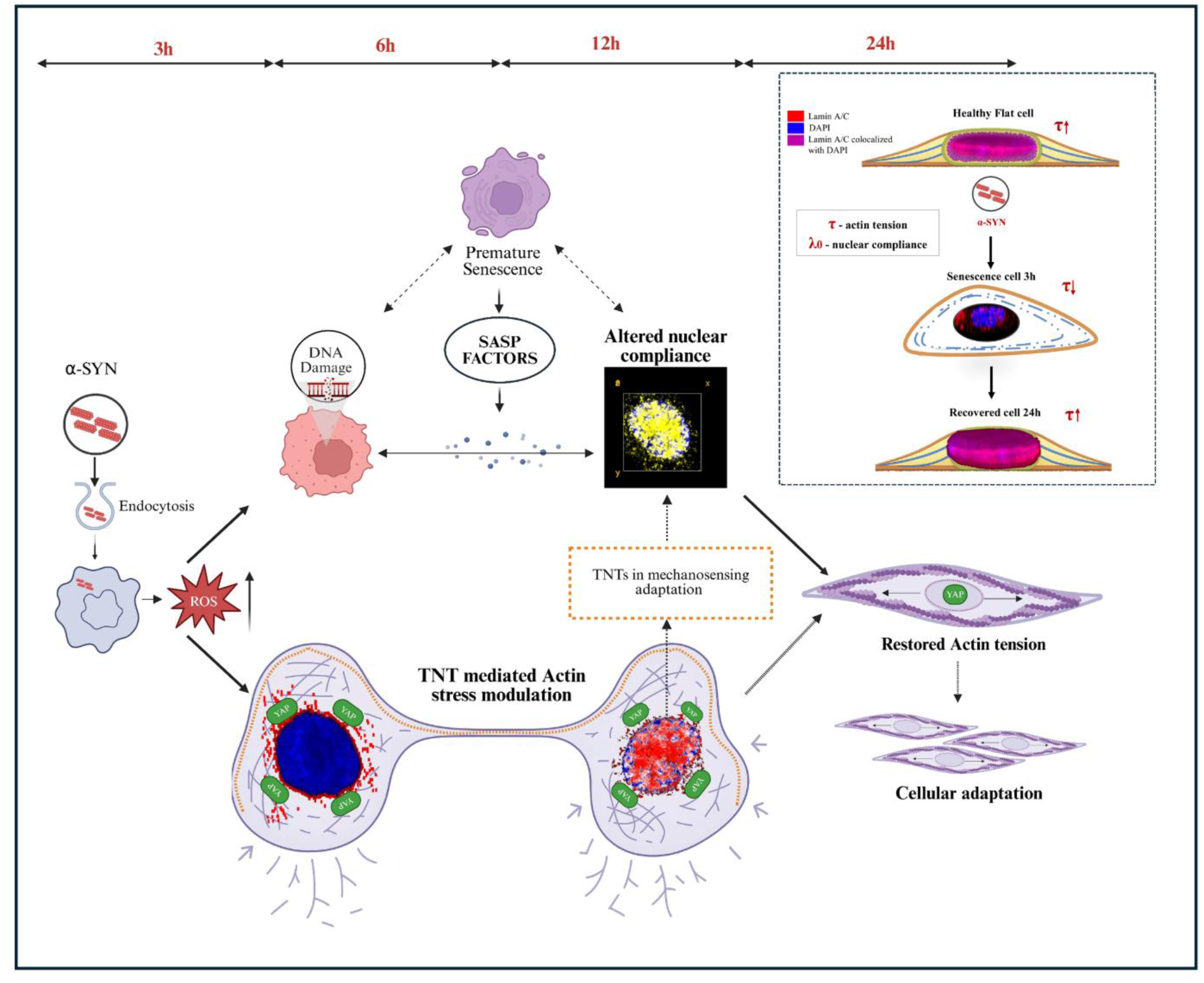

## Introduction

Misfolded pathogenic protein aggregates lead to cell death via proteotoxicity-induced cellular senescence and inflammation. Senescent cells are found in the brain’s glial and neuronal tissues in diseases like Alzheimer’s (AD) and Parkinson’s (PD) (Millett *et al*, 2025). Recently, studies have shown that preformed α-SYN protofibrils induce cellular senescence in astrocytes and microglia in a PD model in a time-dependent manner (Jurk *et al*, 2012). Cellular senescence refers to a perpetual event or state where the cells experience growth arrest due to various stresses (Jurk *et al*, 2012). α-SYN-induced toxicities impair DNA double-strand break repair, resulting in elevated levels of senescence markers (Millett *et al*, 2025). The study showed that deletion of the p21 gene, a crucial signal transducer in cellular senescence, reversed the premature senescent-like traits in the younger mouse model (Verma *et al*, 2021). Cellular stresses, such as oxidative stress, DNA damage, and oncogene activation, can induce premature cellular senescence, independent of irreversible replicative senescence by telomer-end shortening. Stress-induced senescence involving pathways p21 and p53 can be reversed under certain molecular conditions, especially by inducing p53 degradation (Beauséjour *et al*, 2003).

Neurodegenerative aggregates like α-SYN, amyloid-β, and tau cause lysosomal buildup and mitochondrial damage, leading to oxidative stress-induced senescence and the formation of membrane nanotubes or tunneling nanotubes (TNTs) (Dilna et al, 2021; Dilsizoglu Senol et al, 2021). Recent studies have revealed that glial cells, such as microglia and astroglia, play a crucial role in clearing α-SYN aggregates and toxic organelles through TNT-mediated transfer, thereby countering premature senescence and promoting cell survival (Raghavan et al, 2024; Scheiblich et al, 2021; Chakraborty et al, 2023; Rostami et al, 2017a). TNTs are nano-sized, open-ended, actin-rich membrane channels connecting neighboring cells, enabling the transfer of organelles and cytoplasmic materials (Sartori-Rupp *et al*, 2019). The recent visualization of TNTs in brain tissue neurons provides compelling evidence of their relevance to pathophysiology (Chang *et al*, 2025). Cytoskeletal remodeling, especially via actin signaling, is key to TNT formation (Dilna et al, 2021;Raghavan et al, 2024). The elastic tubular structure of TNT facilitates mechanical stress transmission between distant connected cells (Wang *et al*, 2024). The study showed that stretching TNTs by separating two cells modulates actin tension, reducing cytoskeleton curvature.

Senescent cells display reduced cytoskeletal tension via RhoA/ROCK/myosin-II based inhibition, causing nuclear deformations in cancer cells (Aifuwa et al, 2023). The nuclear shape is closely linked to cytoskeletal elements through the linker of nucleoskeleton and cytoskeleton complexes. Therefore, it can be affected by mechanotransduction changes due to the changes in intracellular and extracellular forces (Aifuwa et al, 2023;Kim et al, 2017). The decrease in cytoskeletal tension results in reduced cellular contractility and adhesion, as well as increased mechanical compliance, partly mediated by Lamin A/C. Lamin A/C is an essential structural component of the nuclear lamina, providing mechanical strength to the nucleus and helping to couple it with the cell’s cytoskeleton, thereby forming a critical mechanical link (Lee *et al*, 2007). The cytoskeleton cortex, cell-to-cell junctions, and focal adhesions sense mechanical tension in response to biochemical changes and transmit these signals to the Hippo signaling pathway, which aids cellular homeostasis (Rausch & Hansen, 2020). Defective outside-in mechanotransduction in senescent cells triggers a series of biophysical and biomolecular changes, beginning at the plasma membrane actomyosin cortex, which links the cytoplasm to the nuclear lamina (Aifuwa et al, 2023). A recent study indicates that α-syn protofibrils-treated senescent astroglia promote the formation of TNTs, involving actin modulation via ROCK2 inhibition and nuclear translocation of pFAK (phosphorylated focal adhesion kinase) (Akhter *et al*, 2024). FAK interacts with the Rho-ROCK pathway to modulate the cytoskeleton, which senses mechanical signals from chemical changes or substrate stiffness (Akhter *et al*, 2024). FAK is displaced from adhesion sites in non-adherent cells, reducing actin tension. Rho-mediated activation of ROCK kinases then helps maintain cytoskeletal tension and supports mechanotransduction (Pirone *et al*, 2006). Nucleus size and shape are influenced by cytoskeletal tension and the elastic modulus of the nuclear envelope (Balakrishnan *et al*, 2021). Through a convergent analysis involving mechanical modelling and shape variability studies, the study showed that nucleus shape can be described using two nondimensional parameters, flatness index and scale factor, which are surrogate measures for cytoskeletal tension and nucleus compliance, respectively (Balakrishnan *et al*, 2021). The flatness index positively correlates with actin tension, while the isometric scale factor negatively correlates with the nuclear envelope’s elastic modulus. However, the role of mechanical signaling in TNT biogenesis, particularly how actin cytoskeleton stress influences it and correlates with nuclear lamina integrity in senescence cells, remains unexplored.

This study revealed that reduced actin tension drives TNT formation in astroglia cells (U87 MG astrocytoma cells and primary astrocytes) treated with α-SYN protofibrils, thereby aiding in restoring Lamin A/C integrity and reversing senescence. We used the nucleus flatness index and isometric scale factor from quantitative microscopy to assess this. Small molecules such as Cytochalasin-D, Nocodazole, and Jasplakinolide disrupt TNT formation by affecting cytoskeletal stress through pathways other than ROCK signaling and fail to reverse senescence. STRING analysis of RNA sequencing data from pathway enrichment elucidates the mechanism by which the FAK/Rho/ROCK inhibitory axis governs TNT formation and contributes to mechanotransduction via the Hippo signaling pathway. This process modulates actin cytoskeleton stress and Lamin A/C-mediated nuclear structural integrity, facilitating the survival of α-SYN-protofibrils treated toxic cells by reversing senescence.

## Methods and materials

### Mice

Six to eight-week-old C57BL/6J mice of either sex were sourced from Jackson Laboratories and housed under a 12-hour dark-light cycle at 50–60% humidity and 25°C. Both sexes were equally represented.

### Primary astrocyte culture

C57BL/6J mice were euthanized with CO₂, and their cerebral cortices dissected. The tissues were washed with cold HBSS, and the meningeal layers were removed using sterile Whatman filter paper. Tissues were triturated using a digestion buffer containing 0.25 mg/mL 0.25% FBS (Gibco, #1600004), collagenase (Sigma, #C9409), and 0.25% trypsin-EDTA (Gibco, #25200-072) in 1x PBS with a 1 ml pipette and incubated at 37°C for 30 min. Then the tissue suspension was centrifuged at 1500 rpm for 5 min, the pellet was resuspended, and the cells were plated in MEM with 20% FBS in Corning T25 dishes. The cultured astrocytes were grown and maintained for 14 days before reseeding for the experiments.

### Cell culture

U-87MG (astrocytoma-glioblastoma origin cancer cell lines) cell line was procured from the National Centre for Cell Science (NCCS) cell repository, India. These cells were also tested for mycoplasma contamination. The cells culture were mentained in DMEM (Gibco #2120395) media supplemented with 10% FBS (fetal bovine serum; Gibco #1600004, US Origin), along with 1% PSN (Penicillin-Streptomycin-Neomycin Mixture; Thermo Fisher Scientific #15640055) incubated at 37°C, 5% CO_2_.

### Preparation of α-SYN protofibrils

The human α-SYN wild-type (Addgene ID #36046) construct was purchased and overexpressed in E. coli. Purification was performed using the periplasmic fraction (Raghavan *et al*, 2024) and the samples were aliquoted, flash-frozen in liquid nitrogen, and immediately stored at −80°C. Following its reconstitution with TBS (Tris-buffered saline) and HNE (4-Hydroxy-2-nonenal #Sigma-Aldrich 393204), α-SYN was subjected for seven days to produce protofibrils. Protofibrils were characterised using TEM (Transmission Electron Microscope), as described in our previous study (Raghavan *et al*, 2024).

### Immunocytochemistry (ICC)

After subjecting the cells to the α-SYN protofibrils at the specified concentration (1µM) and time points (3, 6, 12, and 24 hours) along with an untreated vehicle control, the cells were fixed using 4% paraformaldehyde (PFA) and rinsed with 1X phosphate-buffered saline (PBS). The incubation buffer, containing 1 mg/ml saponin and 5% fetal bovine serum (FBS), was applied for 20 minutes at room temperature. Following this step, the appropriate Primary antibodies were applied, and the cells were incubated overnight at 4°C. Afterwards, the cells were washed three times with 1X PBS.

Each secondary antibody, diluted to 1:700, was incubated for 2 hours at room temperature in the dark, followed by additional washes with 1X PBS. The coverslips were mounted onto glass slides with ProLong Gold antifade reagent containing DAPI (Invitrogen P36941). Glass-bottom 35 mm dishes (Cellvis, D35-14-1.5-N) were employed for certain experiments.

The primary antibody Lamin A/C (CST ID #2032S) effectively stains the nuclear envelope. Phalloidin conjugated with iFluor 555 (Abcam #176756) is utilized for staining TNTs. Paxillin (CST # 50195S) Other primary antibodies like p21 (CST # 2947S), p53(Invitrogen # MA5-12557), YAP (Invitrogrn # PA1-46189) and LATS1/2 (Invitrogen # PA5-115498) have been used along with the following secondary antibodies - Alexa 488 goat anti-rabbit (A-11070; Invitrogen), Alexa 555 anti-mouse (A-1413312; Invitrogen), and Alexa 488 anti-mouse IgG (H+L) (A-11059; Invitrogen). All primary antibodies were applied at a dilution of 1:500, secondary antibodies at 1:1000, and phalloidin at 1:700. The immunocytochemically stained cells were imaged using a confocal microscope (Zeiss LSM880, Carl Zeiss, Germany and Olympus FV3000) or a fluorescence microscope (IX73-Olympus).

### Inhibitor treatment

U87MG cells were treated with small molecules such as 5 μM ROCK (Y-27632 # Sigma-Aldrich Y0503), 0.5μM Cytochalasin-D (Thermo Fisher Scientific # PHZ1063), 100nM Nocodazole (Sigma-Aldrich # M1404), and 150nM Jasplakinolide (Invitrogen # J7473), 10mM N-acetylcysteine (Nac) (Sigma # A9165-5G) before being subjected to further experiments.

### Western Blot Quantification

U-87 MG cells were treated with 1µM α-SYN protofibrils for 3, 6, 12 and 24 hours along with an untreated vehicle control, after being seeded at a density of one million per well in a six-well plate. After discarding the post-treatment medium, 1X PBS was used to wash the cells. After adding RIPA buffer, the cells were scraped and collected into 1.5 ml tubes. The tubes were placed on ice and vortexed three times at irregular intervals for fifteen minutes. The suspension was then spun for 10 minutes at 12,000 rpm, and the supernatant was collected and kept at −20 °C. Following the normalisation of the protein content by Bradford reagent (Sigma #B6916), western blot was performed.

Primary antibodies, Actin (PAS097Hu01 Cloud clone dilution 1:1000), P^53^(MAA928Hu22 cloud clone 1:500), P^16INK4a^/CDKN2A (A11651 ABclonal 1:1000), Lamin B1(119D5-F1, Invitogen) (1:1000), 14-3-3ζ (PA5-27317, Invitrogen 1:1000), LATS1/2– (BS-4081R, Invitrogen 1:1000) and Beta Tubulin (32-2600, Invitrogen, 1:1000) were diluted in 5% BSA, after 1h of blocking with 3% BSA. Secondary antibodies, Goat anti-mouse (H+L) (Invitrogen 32430 dilution 1:1500) and Goat Anti-Rabbit (H+L) (Invitrogen 32460 1:1500) were subjected to the blots. The blot was developed with ECL solution (SuperSignal West Femto Trial kit Invitrogen 34094) and was quantified by densitometry analysis using a Fiji software gel analyzer plugin.

### β- galactosidase activity assay

U87MG cells were meticulously seeded at a density of 10,000 cells per well in a 24-well plate to establish optimal growth conditions. Subsequently, the cells were treated with 1 μM α-SYN protofibrils for specified time intervals of 3, 6, 12, and 24 hours, along with an untreated vehicle control. These time points were selected based on preceding experimental data that indicated significant impacts on cellular morphology and viability. Upon the completion of the treatment period, the activity of β-galactosidase was evaluated utilizing a specialized β-galactosidase staining kit (AKR-100, Cell Biolabs Inc), which facilitates the quantification of enzymatic activity in a visually discernible manner. Brightfield images of the stained cells were captured using a color camera, providing a comprehensive visual representation of the staining process and enabling further analysis of the cellular alterations in response to the treatment.

### TUNEL Assay

To measure both the early and late stages of apoptosis, the terminal deoxynucleotidyl transferase-mediated dUTP nick-end labelling (TUNEL) assay was utilised. The The TUNEL assay identifies DNA breakage by incorporating BrdUTP into the free 3’-hydroxyl ends. An anti-BrdU monoclonal antibody labeled with Alexa Fluor 488 is used to detect BrdU. Upon α-SYN protofibrils treatment for durations of 3h, 6h, 12h and 24h, along with vehicle control, U87MG cells were seeded in a 48-well plate and subjected to a TUNEL Assay (Invitrogen Apoptosis BrdU TUNEL Assay kit # A23210) to assess DNA fragmentation following α-SYN protofibrils treatment.

### RNA sequence analysis

RNA was purified, and sequencing was done from U-87 MG cells treated with α-SYN protofibrils for durations of 3h and 24h, along with untreated vehicle controls. The analysis commenced with quality assessment and trimming of raw sequencing reads to ensure high-fidelity data. The cleaned reads were then aligned to the reference genome using a splice-aware aligner, and gene-level read counts were subsequently obtained. To account for technical variability, the raw counts were normalized to correct for differences in sequencing depth and RNA composition. Differential gene expression (DEGs) was evaluated by fitting the normalized counts to a Negative Binomial statistical model, and significance was determined using the Wald test. To prevent false positives from multiple hypothesis tests, p-values were corrected using the Benjamini–Hochberg method to preserve the False Discovery Rate (FDR). Heatmaps for RNA-sequence pathways were created using selected genes by analyzing DEGs (differentially expressed genes) data with EnrichGO software. A few genes in the pathway heatmaps were manually chosen based on their known relevance reported in the literature. The genes were selected from significantly expressed 3-hour α-SYN protofibrils-treated cells compared to control cells based on two criteria: a fold change (FC) ≥ 1 and a false discovery rate (FDR) ≤ 0.05. For the comparison between the 3-hour and 24-hour time points, DEGs were identified considering reverse change in FC, focusing on the same genes that reverted to their baseline regulatory states compared to the 3-hour-treated cells. The majority of the identified genes adhered to the FDR ≤ 0.05, with only a limited number of exceptions. Finally, the biological relevance of the identified DEGs was explored through Gene Ontology (GO) and pathway enrichment analyses. The outcomes of these analyses were visualized using hierarchical clustering heatmaps to highlight key patterns of transcriptional regulation.

### String analysis for protein interaction

**“**The Search Tool for the Retrieval of Interacting Genes/Proteins”, or STRING, is a database that has been designed to gather, analyze, and provide valuable insights into the interactions between proteins, making it a comprehensive and commonly used resource. This resource provides a comprehensive collection of data on physical interactions, like direct protein binding, and functional associations, which describe how proteins may collaborate in biological processes. We utilized STRING analysis (Szklarczyk *et al*, 2023) from the genes obtained through our RNA sequencing data to examine the interactive visualizations of protein networks, predictive interaction scoring systems, and pathway analyses.

### Confocal Microscopy

Confocal laser scanning microscopy (CLSM) was used to capture z-stack images, with high-contrast visuals, using either a Zeiss LSM880 confocal and Olympus FV3000 laser scanning microscope or an Olympus fluorescence microscope. Confocal images were taken using specific objectives and filter sets (DAPI, FITC, TRITC) with 405 nm, 488 nm, and 561 nm lasers. Images were captured with a pixel dwell time of 1.02 μs and an xy-pixel size of 220 nm2. Multiple images (5-10) per condition were taken from random areas. This technique allows for detailed analysis of cytoskeletal dynamics and cellular projections. We generated 3D representations using the 3D Volume View tool in ImageJ, Fiji software, to visualize cellular morphology and spatial relationships. The confocal images were analyzed with Fiji, a Java-based image processing software developed by the NIH and LOCI.

### Volumetric Nuclear Imaging

Nuclear image stacks were obtained using the Olympus FV3000 laser-scanning microscope. A 60x, oil immersion, 1.4 NA objective was used at a volumetric pixel resolution of 0.207×0.207×0.5 μm. The geometric properties of the nucleus were acquired by using a custom MATLAB image processing routine. The shape parameters τ and λ_0_ were determined by fitting the nucleus two-parameter model to the nucleus morphology to a nucleus shape model.

### Quantification of TNT numbers

The quantification of tunneling nanotube (TNT) structures was conducted through an analysis of the z planes. It is important to note that TNTs (less than 1μM in diameter) do not adhere to the substratum; rather, they are located within the intermediate z-stacks. This unique positioning facilitates a comprehensive assessment of their structural characteristics (Valappil *et al*, 2022). The reconstruction of z-stack images was accomplished utilizing the 3D volume view plugin of Fiji. TNTs were manually quantified and represented as a ratio of the number of TNTs to the number of cells within each field.

### Quantification of Lamin Intensity

Lamin A/C expression was analysed by quantifying the labelled protein intensities per cell from images for each condition. Intensities were measured by drawing and selecting regions of interest using the ROI plugin in Fiji.

### Colocalization Analysis

The confocal images were utilized to evaluate the percentage of colocalization. The expression of two proteins in the same cells was analyzed using a color threshold tool in Fiji. The percentage of colocalization or internalization was subsequently calculated based on the area of overlap identified by this tool.

### Superresolution microscopy

Super-resolution advanced Nikon Spatial Array Confocal system (NSPARC). Image stacks were obtained using the Nikon AX laser-scanning microscope. A 40x objective was used at a volumetric pixel resolution of 111μm×111μm and optical resolution of 0.21μm.

### Live Imaging

U87MG cells were seeded at a density of 10,000 cells per well on glass bottom dish. Cells were treated with either 1µM α-SYN protofibrils or 5µM ROCK inhibitor. The glass-bottom dish was precisely positioned within the incubator chamber of the confocal microscope (Olympus FV3000). A diluted solution of CellMask™ Green plasma membrane (PM) stain (37608, Invitrogen) and Hoechst stain was introduced to the media inside the incubator chamber, ensuring that the plates remained undisturbed. Subsequently, live images were taken for analysis.

### Nondimensional mechanical model for the nucleus

The shape of the nucleus is typically characterized by geometrical characteristics, including area, volume, height, and eccentricity. These parameters are generic geometric descriptors and are not unique to the nucleus or its particular physical environment. Hence, there is a need for shape parameters tailored to the nucleus, which we have derived using a mechanical model. Nucleus shape is determined by the mechanical equilibrium between the stress created in the nuclear envelope and the forces acting on the nucleus. Therefore, we hypothesized that, with the use of a suitable mechanical model, the forces and the elastic characteristics of the nuclear envelope could be deduced from the shape. To this end, based on previous experimental studies (Kim *et al*, 2015), we modeled the nucleus as an inflated membrane compressed between two rigid flat plates. The membrane mentioned here refers to the mechanical notion of a membrane. This implies a 2D element that does not resist bending and only develops strain energy due to stretching. In the actual nucleus, this membrane represents the nuclear envelope, which comprises the nuclear membrane and the underlying lamina. The nuclear envelope is modeled as a hyperelastic material (incompressible Mooney-Rivlin material model) and spherical in the unloaded state. By analysing this mechanical configuration using membrane mechanics, we characterized the nucleus shape into two non-dimensional parameters - (i) λ_0,_ which represents isometric scaling, and (ii) τ, which represents flattening. An independent method, variability analysis of nucleus shapes, converged to the same two parameters, further underscoring their importance. Using a variety of studies, involving multiple cell lines, that included growing cells on substrates with different elastic modulus and depolarizing actin filaments and microtubules, we demonstrated that τ correlates with actin tension and λ_0_ correlates with nucleus compliance (Balakrishnan *et al*, 2021; Balakrishnan, 2022).

Here, we used these parameters to deduce alterations in actin tension and nucleus compliance due to various perturbations such as α-SYN protofibrils, Y27632 and cytoskeletal modulators. We treated U87-MG cells with these modulators for 3, 6, 12, and 24 hours and obtained the nucleus shape using confocal microscopy. By fitting the mechanical model to these nucleus shapes, λ_0_ and τ were estimated, which were further used to infer the changes in actin tension and nucleus compliance.

#### Plotting and statistical analysis

All the graphs are plotted in GraphPad Prism software and MATLAB. To assess the significance of the analyzed data, two-way ANOVA or paired ANOVA tests were conducted based on the experimental parameters detailed in the figure legends. The statistics were derived by pooling the data from three biological replicates.

## Results

### TNT-formation correlates with α-SYN protofibrils-induced senescence

In our previous study, we showed that α-SYN protofibrils-induced oxidative stress elevates ROS levels and facilitates TNT formation (Raghavan *et al*, 2024). This leads to premature cellular senescence-like characteristics in primary astrocytes and malignant astrocytoma cell lines (U87 MG and U251), for an initial transient time period (3-6 hours). In this study, we systematically characterized the signaling axis for senescence in U87 MG cells. We observed that the transient formation of TNTs at 3 and 6 hours (Figure 1A) following α-SYN protofibrils treatment correlates with a temporary upregulation of the senescence marker p21 in the nuclei of the cells (Figure 1B). To effectively illustrate cellular senescence, β-galactosidase staining was conducted after treatment with α-SYN protofibrils (Figure 1C). The results identified senescence-positive cells (blue cells) transiently at early time points (3 and 6 hours) (Figure 1C), reinforcing the validity of our observations. Quantification of TNTs (Figure 1D), p21 nuclear expression (Figure 1E), and β-galactosidase (Figure 1F) shows that TNT biogenesis correlates with early α-SYN protofibrils-induced transient premature senescence. Further, a decrease in LaminB1 expression at 3 and 6 hours indicates that the cells are entering a temporary senescent state (Figure 1G, H). After 12 hours, the cells begin to recover from senescence, and TNTs gradually disappear. To assess the reversible nature of senescence, we conducted western blot analysis for p53 (Figure 1I, J), and p16 (Figure 1K, L). The results showed that levels of p53 were found to be elevated during these initial time points (3 and 6 hours) (Figure 1I, J). Notably, the expression of p16 remained consistent throughout the study (Figure 1K, L), indicating that the senescence observed is also a reversible, transient process, similar to the α-SYN protofibrils-induced TNT formation.

**Figure 1:**
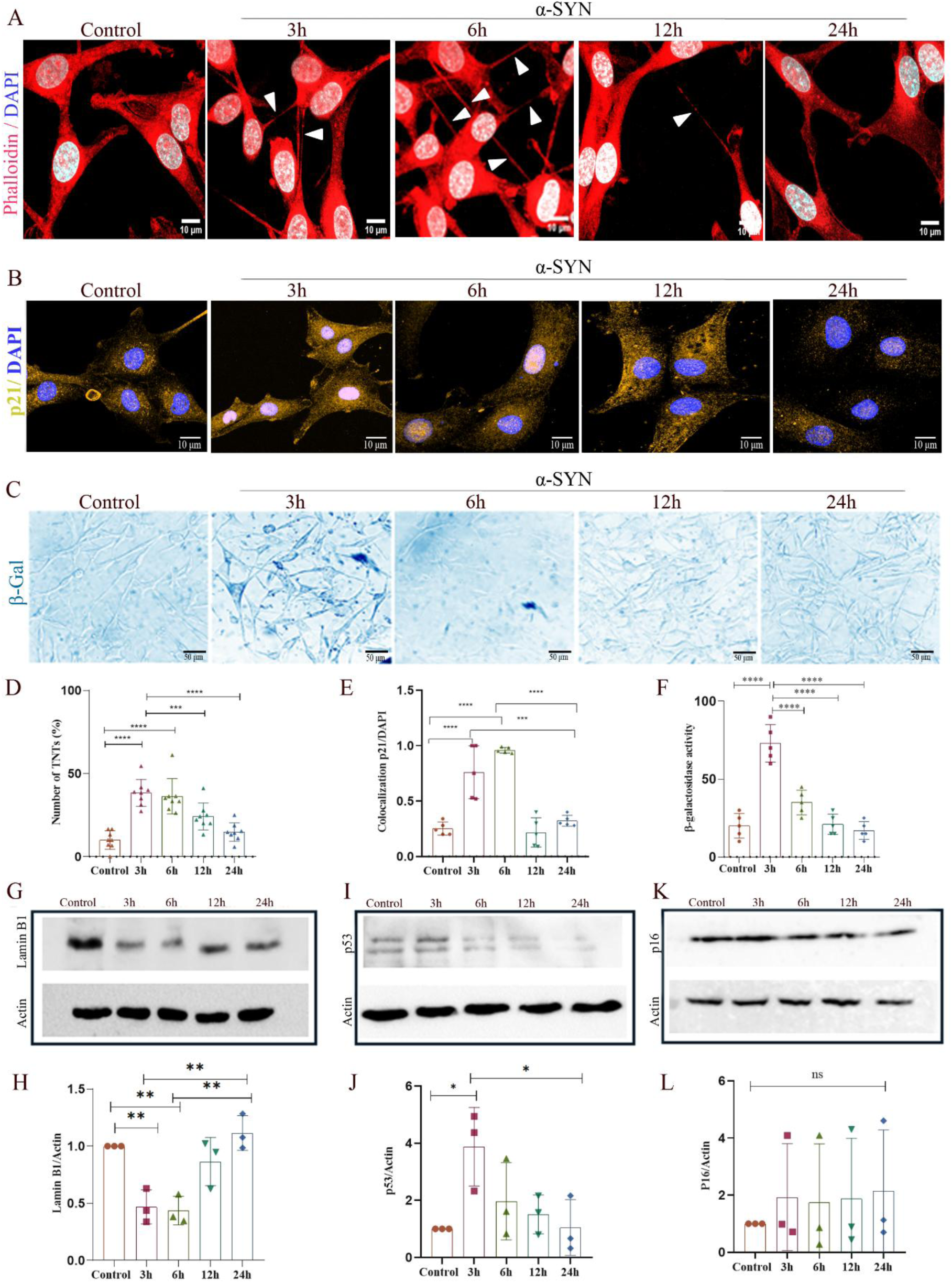
α-SYN-induced cellular senescence correlates with TNT-formation. U87MG cells were treated with α-SYN (1μM) at intervals of 3, 6, 12, and 24 hours. A) Confocal images of phalloidin (red) stained TNTs, B) Nuclear localization of senescence marker p21 and, C) Images of β-galactosidase-stained cells in the α-SYN-treated cells, compared to control cells. The graph’s datapoints represent the number of images used for analysis, pooling data from three biological repeats. Quantification of D) percentage of TNTs by counting the numbers normalized with the number of cells, E) nuclear localization of p21, and F) β-Galactosidase activity analyzing the average intensities of positively stained cells. Western blots of G) Lamin B1, I) p53, and K) p16. Quantifications of H) Lamin B1, J) p53, and L) p16 western blots. Full blots are represented in Supplementary Material 2. Scale bars are denoted on the images. Data are expressed as mean ± SD, ∗∗∗p ≤ 0.001. Statistics were analyzed using a two-way ANOVA. N=3.

### Crosstalk between Lamin A/C and actin tension in α-SYN protofibrils-induced senescent cells

α-SYN protofibrils-treated senescence cells induce nuclear fragility during an initial treatment period of 3-6 hours in U87 MG astrocytoma cells (Raghavan *et al*, 2024). It is well established that the shape of nuclei is influenced by cytoskeletal forces through complexes that link the nucleoskeleton and the cytoskeleton, making the nucleus responsive to mechanotransduction (Lee *et al*, 2007). Reduced cytoskeletal tension can decrease contractility and enhance nuclear compliance through Lamin A/C, a crucial nuclear lamina protein. Thus, in this study, we aim to examine the correlation between actin cytoskeleton changes and nuclear shape following α-SYN protofibrils treatment. Specifically, we intend to investigate how the biogenesis of TNTs may influence the organization and distribution of actin filaments, as well as the consequent effects on the mechanical properties of the nucleus.

We investigated the change in laminA/C integrity (Figure 2A-B) in relation to actin tension (Figure 2C-D) upon α-SYN protofibrils-treatment using the nucleus shape model. For this, we measured the nucleus shape at 3, 6, 12 and 24 hours using confocal imaging (Figure 2C). There were substantial changes in the volume, projected, and surface areas (Figure S1A-C). While volume increased up to 6 hours and recovered by 12 hours, projected and surface areas showed an opposite trend. To obtain actin tension, we used the flatness index, τ, derived by fitting the nucleus morphology to the nucleus shape model (Figure 2C,D). The results show a significant reduction in τ at 3 to 6 hours, indicating a decrease in actin tension, followed by recovery from 12 to 24 hours (Figure 2D). Changes in actin tension are known to cause corresponding changes in Lamin A/C (Buxboim *et al*, 2014). Hence, we investigated whether the modulation of actin tension causes alteration in Lamin A/C expression (Figure 2A).

**Figure 2:**
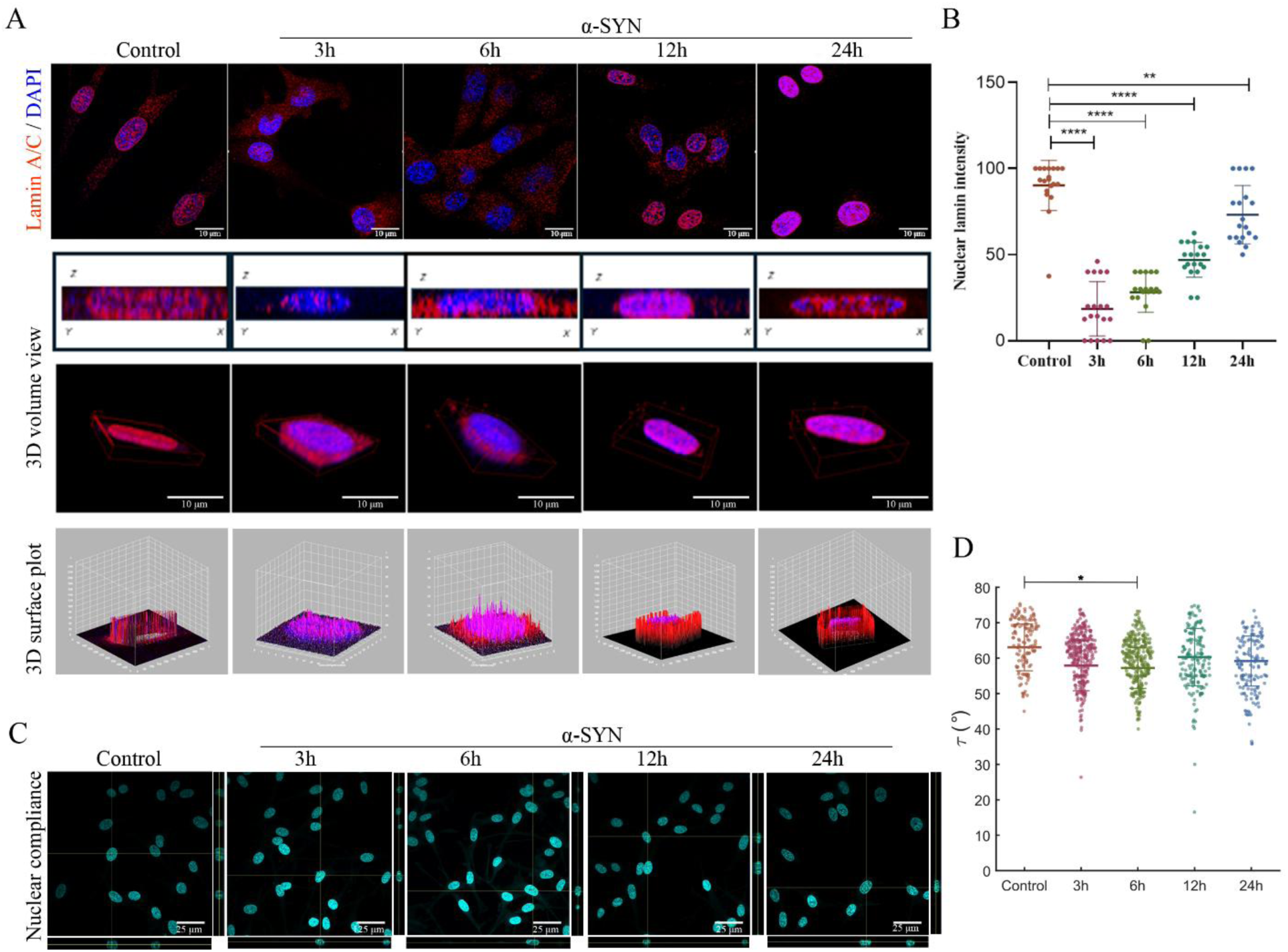
Interplay between Lamin A/C and actin-cytoskeleton tension upon α-SYN treatments. *A)* Fluorescence images of maximum intensity projected confocal z-stacks in U87MG cells stained with Lamin A/C (Red) and DAPI (Blue) upon α-SYN treatment (1μM) for 3, 6, 12, and 24 hours. Represented 3D volume view and 3D surface plots showing Lamin A/C distributions in the nucleus of the α-SYN-treated cell. B) Quantification of Lamin A/C intensity in the nucleus. Data are expressed as mean ± SD, ∗∗∗p ≤ 0.001. Statistics were analyzed using a two-way ANOVA. C) Orthogonal Z-projection of DAPI-stained nuclei in U87MG cells upon α-SYN treatment. D) Quantification of Flattering index (τ) or the actin tension of U87MG cells upon α-SYN treatment. Scale bars are denoted on the images. The graph shows the number of cells used for analysis, pooling data from three biological repeats. Data are expressed as mean ± SD, ∗∗∗p ≤ 0.001. Statistics were analyzed using a paired ANOVA on the means of three independent sets, since data points differ over time points. N=3.

Under typical physiological conditions, the nuclear lamina, which provides essential structural support to propagate mechano-sensing between the nucleus and the cytoskeleton, remains intact. Three-dimensional volume views and surface plots of nuclear Lamin A/C images show that the nuclear lamina surrounding the nucleus remains intact under normal conditions (Movie S1). However, when treated with α-SYN protofibrils, transient deformations of Lamin A/C are observed in the initial hours (3 and 6 hours) (Movie S2). Notably, the staining of the Lamin A/C in the surface returns to normal after 12 and 24 hours of treatment (Figure 2A-B, and Movie S3). Similarly, Lamin B-stained nucleus also showed transient deformation in α-SYN protofibrils-treated cells (Figure S2A-B). Exploring the complex relationship between Lamin A/C and the actin cytoskeleton, this study examines how varying levels of tension within the actin network respond to treatments involving α-SYN protofibrils, leading to the formation of TNTs.

### ROCK inhibition interplays between Lamin A/C and actin-cytoskeleton tension

Previous studies, including ours, have shown that ROCK inhibition facilitates the biogenesis of TNTs (Raghavan et al, 2024, Scheiblich et al, 2021). Further, the Rho-ROCK signaling pathway is crucial for maintaining the equilibrium between Lamin A/C and actin-cytoskeleton tension and mechanotransduction. Hence, we investigated the effect of ROCK inhibitor (Y-27632), on TNTs, actin tension and Lamin-A/C.

Similar to α-SYN protofibrils, we found that inhibition of ROCK pathway using Y-27632 resulted in the dysregulation of Lamin A/C around the nucleus and facilitation of TNT formation (Figure 3A-D). Three-dimensional volume views and surface plots of nuclear lamin images demonstrate deformations of Lamin A/C upon Y-27632 treatment in the U87-MG cells, where nuclear lamina surrounding the nucleus is intact under normal conditions (Figure 3A,C). Quantification of TNT data shows Y-27632 treatment promotes the biogenesis of TNTs over the time of treatment till 24 hours (Figure 3B, D). Actin tension measured using the flatness index, τ, from the nucleus shape model also showed a similar trend, a decrease until 6 hours and then partial recovery from 12 to 24 hours (Figure 3E). Lamin A/C also showed a similar trend to α-SYN protofibrils, a decrease until 6 hours and partial recovery up to 24 hours. These findings suggest a potential role of ROCK-mediated reduced actin stress in the formation and maintenance of TNTs, which may have important implications for intercellular communication and cellular homeostasis.

**Figure 3:**
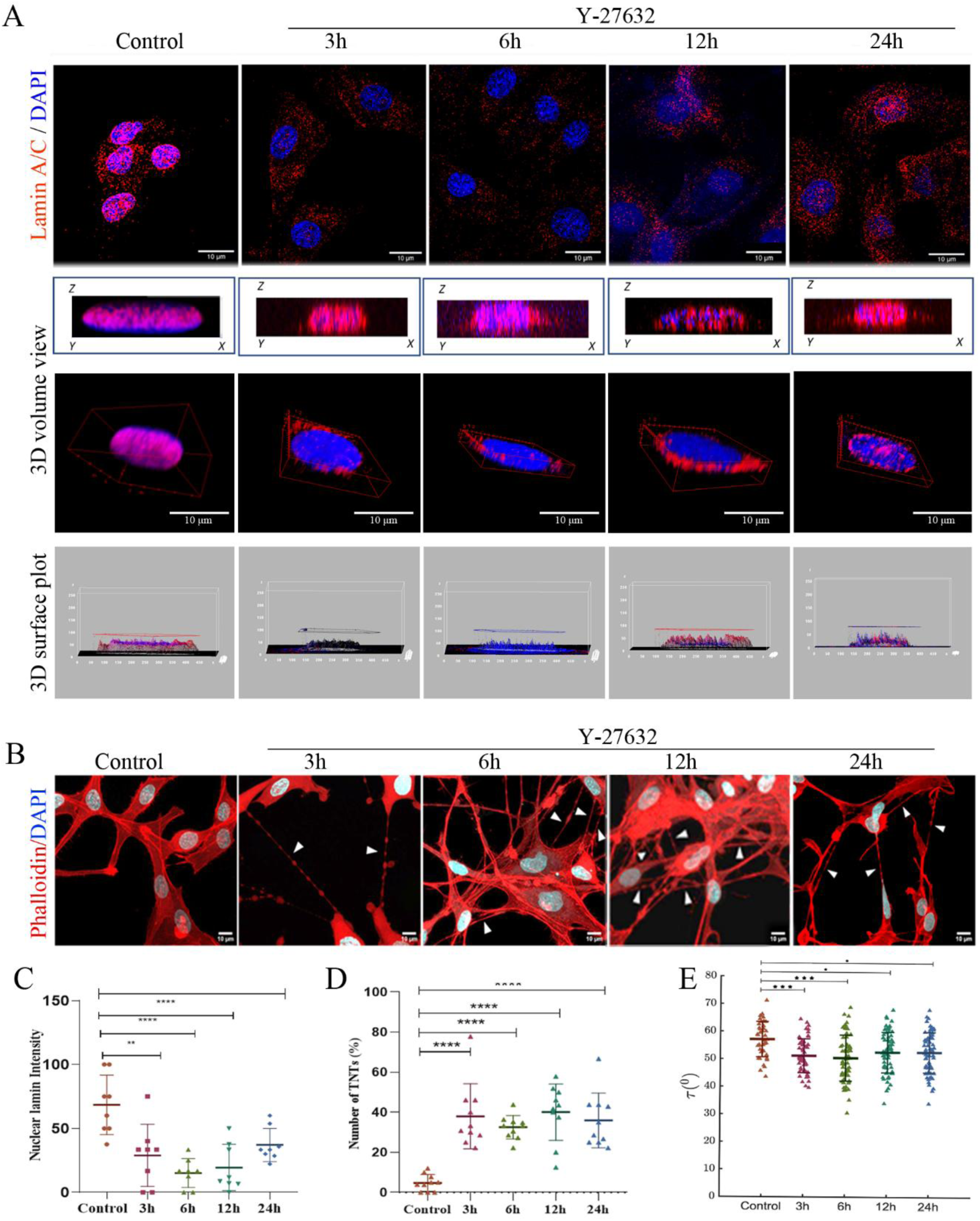
ROCK inhibition interplays between Lamin A/C and actin-cytoskeleton tension. A) Fluorescence images of maximum intensity projected confocal z-stacks of U87MG cells stained with Lamin A/C (red) and DAPI (blue) upon ROCK inhibitor Y-27632 treatment (5μM) for 3, 6, 12, and 24 hours. Represented 3D volume view and 3D surface plots showing Lamin A/C distributions in the nucleus of the Y-27632-treated cell. B) Confocal images of Y-27632-treated cells stained with Phalloidin (red) and DAPI (blue) to visualize the number of TNTs. The graph datapoint represents the number of images used for analysis, pooling data from three biological repeats. C) Quantification of Lamin A/C intensity in the nucleus upon Y-27632 treatment. D) Quantification of percentage of TNTs by counting the numbers normalized with the number of cells treated with Y-27632, and E) Quantification of flattering index (τ) or the actin tension of U87MG cells upon Y-27632 treatment. Scale bars are denoted on the images. Graph datapoints represent the number of cells used for analysis, pooling the data from three biological repeats. Data are expressed as mean ± SD, ∗∗∗p ≤ 0.001. Statistics were analyzed using a two-way ANOVA. N=3.

### Actin tension modulators that inhibit TNTs are ineffective in reversing α-SYN protofibrils-induced senescence

Our results indicate that TNT formation in α-SYN protofibrils-induced senescent astroglia is caused by decreased actin tension, which occurs through inhibition of the Rho-ROCK pathway. Next, we asked whether reduced actin tension, apart from the Rho-ROCK pathway, can alone induce TNTs. We used three small molecules that modulate the actomyosin network via alternative actin pathways that are not regulated by Rho-ROCK (Kretschmer *et al*, 2019), namely, Jasplakinolide (Jasp), Nocodazole (Noco), and Cytochalasin D (Cyto D). While Jasp stabilizes existing actin filaments, forming actin stress fibers, Noco stabilizes these stress fibers by destabilizing microtubules, increasing actin tension, whereas Cyto D reduces actin tension by depolymerizing actin filaments (Zhang *et al*, 2012; Verstraelen *et al*, 2017).

We observe that these molecules fail to restore the structural integrity of Lamin A/C and disrupt TNT formation in α-SYN protofibrils-treated cells (Figure 4 and S3). Lamin A/C degradation and disorganization of the actin meshwork, with numerous TNTs, are evident only in α-SYN protofibrils-treated senescent cells at 3 hours (Figure 4A-C). The α-SYN protofibrils-treated cells regain Lamin A/C integrity and actin meshwork after 24 hours (Figure S3A-B). We observe that Jasp, Noco, and Cyto D fail to restore the disorganized actin cytoskeleton and the integrity of Lamin A/C in α-SYN protofibrils-treated cells, both at 3 and 24 hours (Figure 4A-B, S3A-B). These small molecules inhibit the formation of TNTs in α-SYN protofibrils-treated cells (Figure 4C, S3C). The molecule Nac, which neutralizes cellular ROS, caused a lesser extent of Lamin A/C damage, actin cytoskeleton meshwork disruption, and TNT formation upon α-SYN protofibrils treatment (Figure 4A-C, S3A-C). Cells were exposed solely to these small molecules without α-SYN protofibrils to assess how these actin modulators affect the cells. Results show that Cyto D and Nac do not alter nuclear Lamin A/C, whereas Jasp partially disrupts and Noco significantly alters nuclear Lamin A/C distribution after 3h of treatment with the respective concentrations mentioned in the materials and methods (Figure S4A-B).

**Figure 4:**
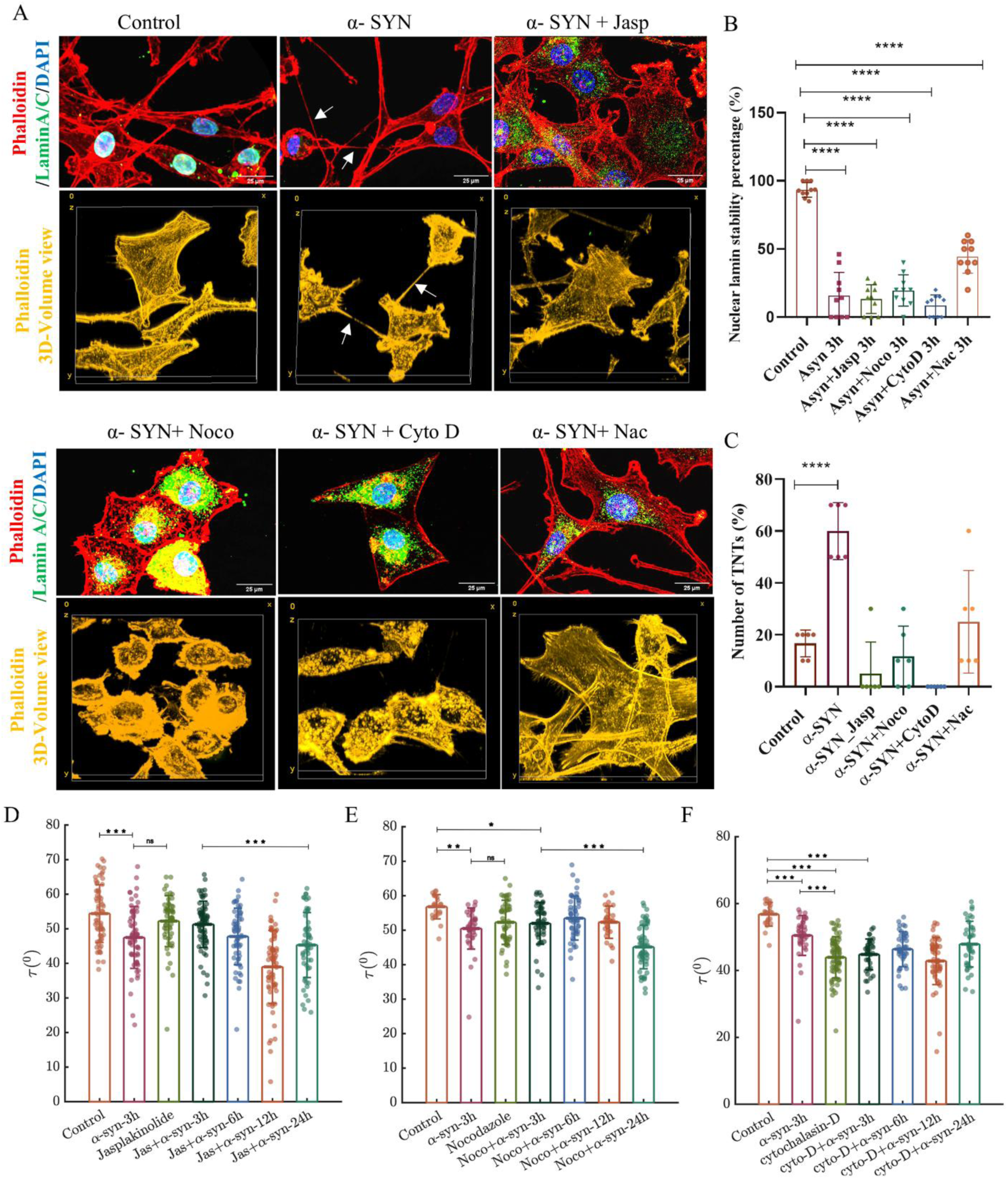
Actin tension-modulating molecules inhibit TNTs and aggravate α-SYN-induced senescence. A) Phalloidin (Red), Lamin A/C (Green) and DAPI (Blue)-stained U87 MG cells treated with α-SYN and α-SYN-treated cells with respective inhibitors Jasp, Noco, Cyto D, and Nac at 3 hours. Phalloidin (yellow) stained 3D volume view of actin cytoskeleton structures obtained from super-resolution microscopy is represented. B) Quantification of Lamin A/C intensities at the nucleus, and C) TNT numbers in percentage, after treatment with the inhibitors in the presence of α-SYN at 3 hours. The graph shows the number of images used for analysis, pooling data from three biological repeats. Quantification of Flattering index (τ) of the cells treated with D) Jasp, E) Noco, and F) Cyto D after treatment of the cells with the inhibitors in the presence of α-SYN for 3,6,12 and 24 hours. The graph shows the number of cells used for analysis, pooling data from three biological repeats. Data are expressed as mean ± SD, ∗∗∗p ≤ 0.001. Statistics were analyzed using a two-way ANOVA. N=3.

Furthermore, we quantified actin stress by measuring nucleus flatness in response to α-SYN protofibrils treatment with Jasp, Noco, and Cyto D, and the results similarly indicate a failure to restore actin stress, consistent with our microscopic observations of actin meshwork restoration (Figure 4D-F). Jasp and Noco, which increase actin-cytoskeleton tension, delay the reduction in actin tension with α-SYN protofibrils treatment at early time points (3 and 6 hours); however, they do not restore tension at later times (12 and 24 hours). Cyto D, which decreases actin-cytoskeleton tension, causes an additional reduction in actin tension with α-SYN protofibrils treatment over time (3-24 hours). Overall, our results indicate that TNT structures are essential for re-establishing Lamin A/C and the actin cytoskeleton tension in α-SYN protofibrils-treated astroglia cells. Small molecules that modulate actin and disrupt TNTs fail to restore Lamin A/C or the actin cytoskeleton network in these cells.

### α-SYN protofibrils alters DEGs and restores p21/p53-mediated oxidative stress and DNA damage pathways

Extracellular α-SYN protofibrils - induces proteotoxic stress, which increases ROS levels and activates the oxidative stress-related pathway (Raghavan *et al*, 2021). Oxidative stress establishes a foundation for a phenotype that resembles premature senescence and contributes to DNA damage (Puspita *et al*, 2017). α-SYN protofibrils promotes TNT formation via ROCK-mediated actin remodulation, enabling cell-to-cell transfers of materials through TNTs in glial cells (Raghavan *et al*, 2024). To clarify these findings, we analyzed the differentially expressed genes (DEGs) from RNA sequencing data of U87 MG cells treated with α-SYN protofibrils for 3 and 24 hours. Notably, several genes involved in the oxidative stress-related cellular senescence pathways were significantly upregulated after 3 hours of treatment, compared to untreated control cells (Figure 5A). After 24 hours, the same genes showed significant downregulation compared to their 3-hour expression (Figure 5A), suggesting that the initial stress response may have alleviated or transformed into another state. The genes IL6, EDN1, IL11, TSP1, DLX2, FHL2, SIK1, and DKK1 trigger p21/p53 pathway-related senescence in response to stress associated with elevated ROS or DNA damage (Figure 5A). Treatment with α-SYN protofibrils initially (at 3 hours) upregulates expression of DNA repair genes (RAD1, HMGB1, CARD8-AS1, CHAC2, FGF18, CENPU, and OVCA2), indicating an early response to oxidative stress-mediated DNA damage. Later (at 24 hours), their expression is downregulated (Figure 5B), indicating that damaged DNA has been repaired. This temporal shift highlights the dynamic nature of gene regulation in response to α-SYN protofibrils treatment, probably orchestrated by TNT formation through actin cytoskeleton remodulation.

**Figure 5:**
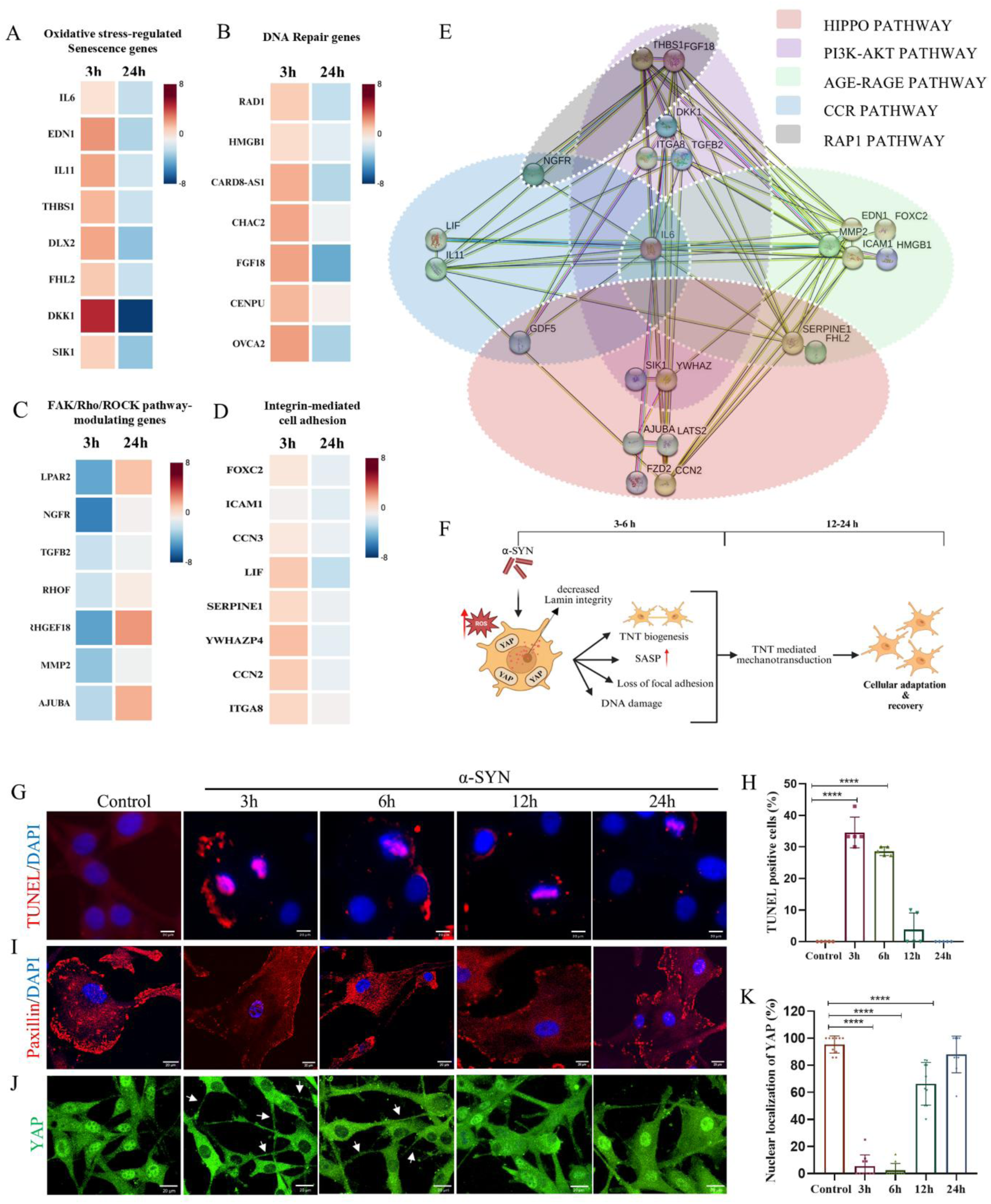
RNA sequencing data reveal mechanical signaling Hippo-pathway in TNT formation upon α-SYN treatment: A) Log2 fold changes of the selected genes from pathway analysis with significant p-values were depicted in the heatmap. This represents the DEGs in U87MG cells after treatment with α-SYN (1 μM) for 3 hours compared to the control, as well as the same genes after 24 hours of treatment compared to the 3-hour mark. The categories include: A) oxidative stress-related senescence genes, B) genes associated with DNA repair pathways, C) genes involved in the FAK-Rho-ROCK signaling pathway, and D) Intrigin-mediated cell adhesion pathway. E) STRING analysis of DEGs maintained in the pathways (A-D). P-value adjusted <0.05. F) Schematic of α-SYN-mediated oxidative stress-induced senescence, highlighting DNA damage, nuclear lamin integrity, and the ROCK-inhibitory pathway that facilitates TNT formation and the reversal of senescent cells. G) Images of TUNNEL assay represent DNA damage, H) quantification of the cells with damaged DNA, I) Paxillin represents FAs, J) cytosolic and nuclear translocation of YAP, and K) quantification of cells with nuclear YAP, in α-SYN (1 μM) treated U87 MG cells for 3,6,12, and 24hours. The graph’s data points represent the number of images used for analysis, pooling data from three biological repeats. Data are expressed as mean ± SD, ∗∗∗p ≤ 0.001. Statistics were analyzed using a two-way ANOVA. N=3.

### α-SYN protofibrils induces TNT biogenesis, modulating FAK/Rho/ROCK and integrin-mediated cell adhesion pathways

In our previous study, we demonstrated that phosphorylated focal adhesion kinase (pFAK) translocates from the cell membrane to the nucleus in senescent astroglia, leading to ROCK2 inhibition, which dynamically modulates the actin cytoskeleton and facilitates the formation of TNTs (Raghavan *et al*, 2024). Similarly, we observe here significant downregulation of FAK/Rho/ROCK pathway-related genes in α-SYN protofibrils-treated senescence cells at 3 hours compared to control cells (Figure 5C). The same set of genes was upregulated again after 24 hours compared to their 3-hour expression, with reversal of senescence toxicities and disappearance of TNTs (Figure 5C). LPAR2 and NGFR downregulation can lead to ROCK inhibition, and the downregulation of RHOF, ARHGEF18, TGFβ2, and AJUBA is directly related to inhibition of FAK activity. MMP2, the key protein that modulates FAK upregulation (Jin *et al*, 2007). LATS1/2 proteins dynamically regulate the FAK-mediated Rho/ROCK pathway by sensing changes in actin cytoskeleton tension, which is downregulated at 3 hours upon α-SYN protofibrils treatment. Further analysis of DEGs data shows that the integrin-mediated cell adhesion pathway (FOXC2, ICAM1, CCN3, LIF, SERPINE1, YWHAZP4, CCN2, ITGA8) is upregulated in 3-hour α-SYN protofibrils-treated senescent cells, and the same genes revert to normal levels at 24 hours. We have observed that non-adherent cells frequently form TNTs with spread, adherent cells, which promote their re-adhesion (Movie S4).

### TNTs in actin stress restoration and mechanical adaptation via Hippo signaling Pathway

Our experimental data aligns with DEG genes, showing that α-SYN protofibrils causes oxidative stress, which leads to a temporary senescence-like state characterized by lamin loss, DNA damage, FAs inhibition, and cell deadhesion in the early hours. TNT formation then modulates the actin cytoskeleton, helping to reverse senescence and promoting adhesion and survival (Figure 5F). Therefore, we searched for interacting protein partners within the pathways mentioned in RNA-sequencing heatmap data in Figure 5A-D, performing a STRING analysis (Figure 5E). STRING analysis shows strong correlations among genes involved in p21/p53-mediated senescence, DNA repair, FAK/Rho/ROCK-related pathway, and the integrin-mediated cell adhesion (Figure 5E). KEGG pathway enrichment suggests significant engagement in biological functions related to the Hippo signaling pathway. The other KEGG enrichment pathways that show significant engagement are AGE-RAGE signaling, PI3K-AKT signaling, cytokine-cytokine receptor (CCR) interaction, and the Rap1 signaling pathway (Supplementary Table S1).

Furthermore, experimental results confirmed the RNA sequence data. The TUNEL assay demonstrated DNA damage, while paxillin staining indicated inhibition of FAs upon α-SYN protofibrils treatment at early time points (3 and 6 hours), followed by DNA repair and cell re-adhesion at later stages (12 and 24 hours) (Figure 5G-I). Finally, the cellular distribution of YAP, a key Hippo-signaling molecule, was observed. Results show that YAP translocates to the cytoplasm to restore mechanical balance when actin tension is low, as seen in α-SYN -protofibrils treated senescence cells at 3 and 6 hours, and at later times, YAP returns to the nucleus (Figure 5J-K). In the canonical Hippo signaling pathway, LATS1/2-mediated phosphorylation of YAP (at Ser127) binds to the cytosolic protein 14-3-3ζ, sequestering YAP in the cytoplasm and inhibiting its nuclear transcriptional activity. Similarly, our WB results show increased levels of LATS1/2 and 14-3-3ζ in α-SYN protofibrils-treated astroglia at initial times 3 and 6h, which correlates with the cytosolic translocation of YAP and senescence (Figure S5)

### TNT reinstates cytoskeletal tension via dynamic mechanical adaptation, followed at single-cell level

TNTs are actin membrane structures, facilitated by the dynamic interplay of actin cytoskeleton under various cellular stress related to oxidative stress. Cells rely on biochemical signals to adapt to fluctuating mechanical stress, a process critical for their survival. To understand the dynamic mechanical adaptation with biochemical changes, we followed single cells over time upon α-SYN protofibrils treatment. We measured the nucleus flatness index and isometric scale factor from quantitative microscopy from confocal z-stacks of the same cell over time and followed the biogenesis of TNTs. The PM of U87-MG cells was labelled with CellMask (green) and the nucleus with Hoechst (blue) to follow the dynamic changes in mechanical stress, marking live cells at the single-cell level upon α-SYN protofibrils treatment (Figure S6A). Single-cell imaging shows significant numbers of TNT-biogenesis upon α-SYN protofibrils treatment for 3 hours. TNT-biogenesis correlates with a decrease in actin stress measured from nuclear flatness (Figure S6B). However, single-cell-based live cell measurements did not show recovery to the same extent as we obtained from the bulk analysis of the fixed cell’s nucleus after α-SYN protofibrils treatment. The sole limitation of single-cell analysis was prolonged exposure to the laser beam up to 24 hours, coupled with the membrane dye, which exhibits toxicity with extended exposure and inhibits the cells from complete recovery. As the cell membrane continued to undergo endocytosis as a normal physiological process, it was necessary to administer the dye at intervals of three hours, which had a toxic effect on the cells. TNTs disappear after 12 hours of α-SYN protofibrils treatment, and some cells regain actin stress after 24 hours (Figure S6A, B). Consequently, while the cells didn’t fully recover in 24 hours, the actin tension indicated some improvement compared to the state after 3 hours. Moreover, the relationship between actin dynamics and nuclear mechanics reveals a sophisticated layer of regulation, suggesting that cellular architecture is not merely a passive structure but an active participant in maintaining homeostasis and responding to biochemical changes.

### TNT Inhibition via Actin Tension Modulators Aggravates p21/p53-Dependent Senescence

Finally, to confirm that TNT formation modulates actin stress in senescent cells and aids recovery, we used small molecules such as Jasp, Noco, and Cyto D. These molecules interfere with TNTs by modifying cytoskeletal stress through alternative actin pathways that bypass the typical Rho-ROCK regulation. The findings indicate that while TNTs are inhibited by these small molecules, they cannot reverse α-SYN protofibrils-induced senescence and cell toxicity mediated by p21/p53. The expression of p21 and p53 remains higher at both 3 hours and 24 hours of α-SYN protofibrils exposure, when treated along with Jasp, Noco, and Cyto D. However, only α-SYN-treated cells show reduced levels of p21 and p53 expression in their nuclei after 24 hours of recovery time (Figure 6A). Nac, that inhibits ROS, partially inhibits p21 expression in the nucleus of α-SYN protofibrils-treated cells at both 3 and 24 hours (Figure 6A). Nuclear expression of p21 and p53 was quantified and shown in the graphs upon α-SYN protofibrils treatment, in the presence and absence of the small molecules Jasp, Noco, Cyto D, and Nac (Figure 6B-C). Overall, the results show that actin stress modulators that prevent TNT formation cannot reverse p21/p53-driven senescence.

**Figure 6:**
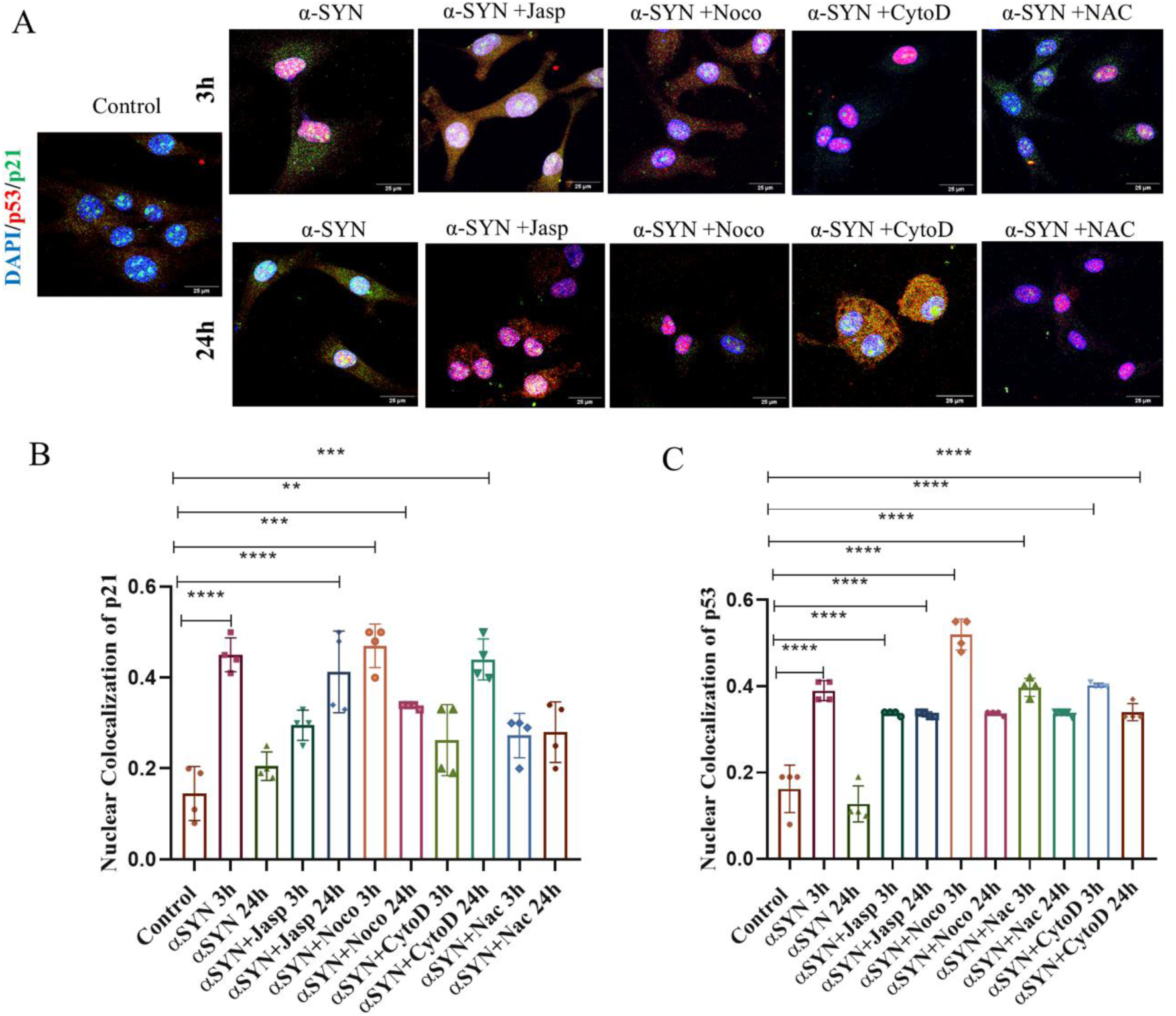
Inhibition of TNTs by molecules that modulate actin tension promotes senescence-related toxicities mediated by p21 and p53. A) Images depict the levels of p21 and p53 expression in the nucleus of α-SYN and α-SYN treated with respective inhibitors Jasp, Noco, Cyto D, and Nac in U87 MG cells at 3 and 24 hours. Quantification of B) p21 and C)p53 levels in the nucleus of the images that are represented in panel (A). The graph’s data points represent the number of images used for analysis, pooling data across three biological repeats. Data are expressed as mean ± SD, ∗∗∗p ≤ 0.001. Statistics were analyzed using a two-way ANOVA. N=3.

### Validation of Mechanosensing signaling-driven TNT biogenesis in primary astrocytes

In a previous study (Raghavan *et al*, 2024), we showed that primary astrocytes develop transient TNTs upon treatment with α-SYN protofibrils, similar to U87-MG cells. Thus, we examined whether astrocytes form TNTs via a similar mechanosensing signaling pathway. Immunostaining data show that YAP translocates to the cytoplasm of astrocytes in α-SYN protofibrils-treated cells at 3 and 6 hours, when TNT formation is observed. Later, at 12 and 24 hours, YAP moves back to the nucleus, and TNTs disappear (Figure 7A-C). YAP, a crucial molecule in Hippo signaling, localizes over TNTs and colocalizes with actin (Figure 7A and Movie S5). Interestingly, we observe that either cell or both cells must contain YAP in the cytosol to form TNTs between them. In contrast, when both neighboring cells have YAP in the nucleus, they do not form TNTs (Figure 7D -E, and Movie S5). We observe many instances in which TNTs form between cells with low actin tension and Hippo-signaling on and those with relatively higher actin tension and Hippo-signaling off (Movie S5). The most notable observation is the high abundance of YAP inside the TNTs, which seems to colocalize with actin, as shown by the surface plot image of α-SYN protofibrils-treated TNT (Figure 7D). α-SYN protofibrils-treated senescent astrocytes also show degraded Lamin A/C at an early time (3 hours) (Figure 7F-G), which correlates with reduced actin stress. The results confirm that cultured primary astrocytes form TNTs by activating Hippo signaling, using a mechanism similar to that in U87-MG astrocytoma cells, to restore mechanical homeostasis when actin tension is low in α-SYN protofibrils-treated senescent cells.

**Figure 7:**
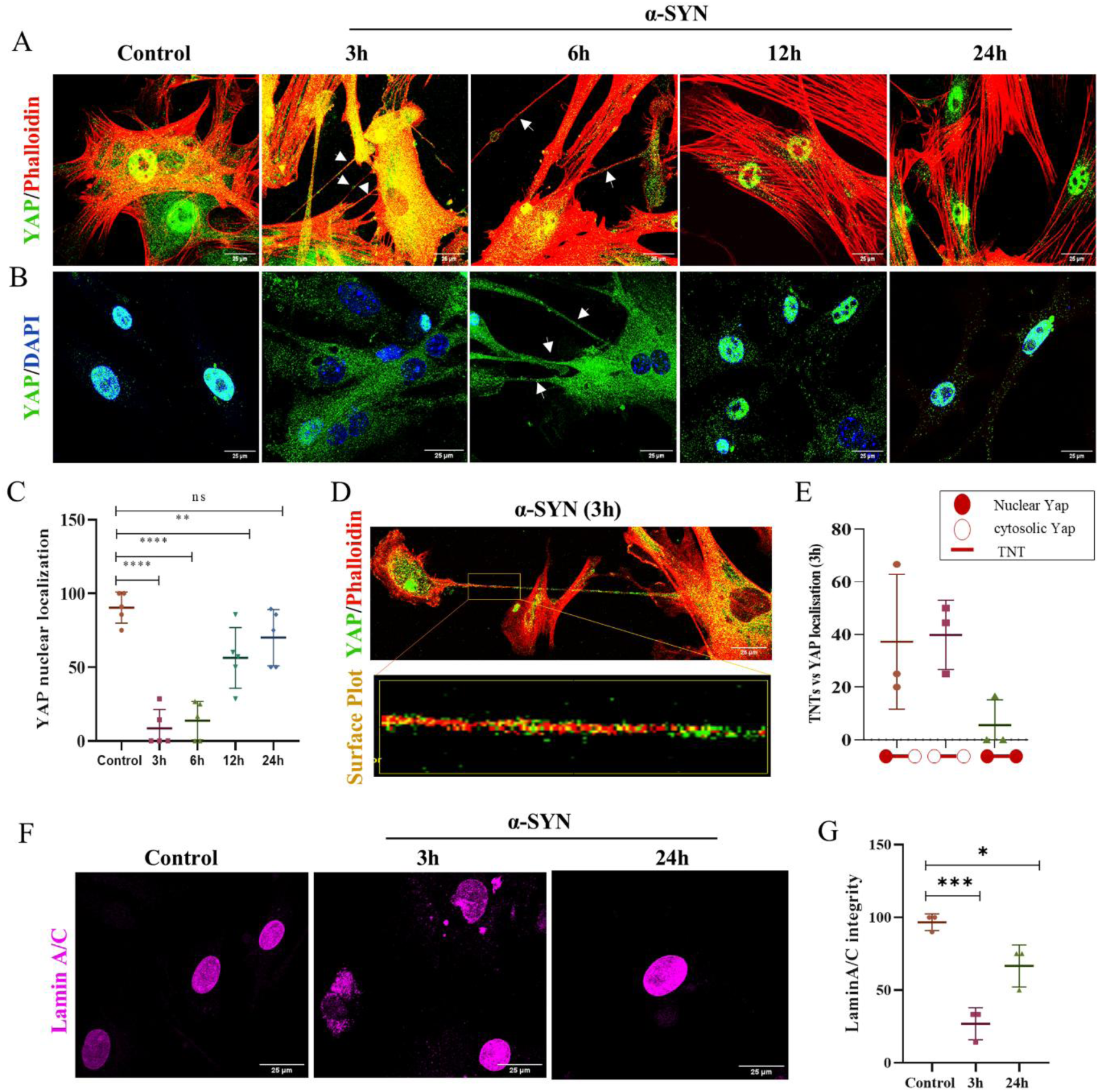
Mechanical signaling Hippo-pathway in TNT formation in astrocytes upon α-SYN treatment: A) Phalloidin (Red), and YAP (Green)-co-stained and B)YAP (green)-stained astrocytes treated with α-SYN for 3 ,6 ,12 and 24 hours. C)quantification of YAP localization in the nucleus. D) YAP (Green) and Phalloidin (Red) co-localization in the TNT. E) Percentage of TNTs between cells: with either cytosolic, and/or nuclear YAP localization. F) Lamin A/C-stained astrocytes treated with α-SYN for 3 and 24 hours. Quantification of nuclear G) Lamin A/C. The graph shows the number of images used for analysis, pooling data across three biological repeats. Data are expressed as mean ± SD, ∗∗∗p ≤ 0.001. Statistics were analyzed using a two-way ANOVA. N=3.

## Discussion

α-SYN aggregation is the most prevalent pathology in PD. The accumulation of α-SYN in brain cells, including neurons and astrocytes, causes oxidative stress and senescence by impairing mitochondria and triggering ROS-induced DNA damage responses (Muwanigwa *et al*, 2024). Recent studies demonstrated that aggregates of α-SYN protofibrils promote the formation of TNTs in astrocytes and astrocytoma cells, and cell-to-cell transfer via TNTs facilitates the clearance of these aggregates and aids cell recovery (Raghavan et al, 2024; Rostami et al, 2017; Rostami et al, 2021; Loria et al, 2017). Previously, we showed that α-SYN protofibrils-induced TNT formation via pFAK-ROCK-mediated actin modulation reverses proteotoxicity-related organelle (lysosome and mitochondria) toxicities, restores cells from premature cellular senescence-like traits, and enhances astroglia survival (Raghavan *et al*, 2024). We observed that exposure to toxic α-SYN protofibrils causes widespread toxicity, with fragmented mitochondria, low membrane potential, and cell-to-cell mitochondrial transfer via TNTs aiding survival. Several studies indicate that healthy mitochondrial transfer between cells is the key mechanism underlying glial survival in response to neurodegenerative proteotoxicity (Scheiblich et al, 2021;Chakraborty et al, 2023). However, the mechanism underlying TNT-mediated survival of the widely spreading, highly toxic astroglia population treated with α-SYN protofibrils cannot be explained solely by cell-to-cell mitochondrial transfer. In this study, we reveal that α-SYN protofibrils-induced senescent astroglia survive by undergoing mechanotransduction that promotes the formation of TNTs. We observe that this process involves a complex interaction between nuclear structural integrity, regulated by Lamin A/C, and the actin cytoskeleton network, which maintains mechanical stability through the Hippo signaling pathway.

We have demonstrated that astrocytoma cells exposed to α-SYN protofibrils undergo a temporary transition into a senescence-like state, characterized by the activation of the p21/p53 pathway. This results in decreased cytoskeletal stress and the temporary formation of TNTs. RNA sequence analysis further shows that actin stress and TNT formation in α-SYN protofibrils-treated senescent cells during early hours (3 and 6 hours) are linked to transient increases in p21/p53 pathway-associated oxidative stress-related senescence markers. Premature reversible senescence is characterized by senescence-associated β-galactosidase activity (SA-β-gal) and a decrease in the levels of Lamin B1. α-SYN protofibrils did not change P16 levels, a marker of replicative senescence that permanently stops cell division (Beauséjour *et al*, 2003).

Degradation of nuclear lamin is a common phenotype in cells that undergo senescence induced by oxidative stress. Nuclear lamin form a peripheral nucleoskeleton and help maintain the nuclear shape, closely connecting to cytoskeletal components. The reduction in cytoskeletal tension results in lower cellular contractility and adhesion, accompanied by increased mechanical compliance, which is partly controlled by lamin proteins (Lamin A/C and B1) (Nava *et al*, 2020). Our results show that α-SYN protofibrils triggers reversible premature senescence at early time points (3 and 6 hours), leading to nuclear lamina breakdown and cell de-adhesion. Using the nucleus flatness index and isometric scale factor from quantitative microscopy, we demonstrate that Lamin A/C plays a crucial role in regulating actin stress in α-SYN protofibrils-treated senescent cells. Cells with Lamin A/C disruption exhibit reduced actin stress in response to α-SYN protofibrils treatment at early time points (3 and 6 hours), thereby promoting TNT formation. A study shows that mutant KRas can transfer between tumor cells via TNTs, reducing membrane tension and increasing phospholipid flow. This change in membrane mechanics also enhances the metastatic and invasive abilities (Zheng *et al*, 2024). TNTs transfer mechanical stimuli between cells and activate mechanosensitive calcium channels to establish actin cytoskeleton tension (Wang *et al*, 2024). TNT formation via actin polymerization may restore actin tension, reverse Lamin A/C loss, and the senescence state.

Mechanotransduction influences chromatin remodeling by detecting lamina-associated heterochromatin under mechanical stress, helping to sustain the nuclear envelope and regulate transcription (Kovacs *et al*, 2023). This process assists in recovering from senescence-related stress, and DNA damage is universally recognized as a key cause of cellular senescence. We observe a transient DNA damage state in astroglia following α-SYN protofibril treatment during early premature cellular senescence. α-SYN protofibrils also plays a role in maintaining genomic stability, with its nuclear pool facilitating DNA repair through DNA binding (Millett *et al*, 2025; Jos *et al*, 2025). Senescent cells also exhibit decreased cytoskeletal tension through the inhibition of RhoA/ROCK/myosin II, resulting in nuclear deformations in cancer cells (Aifuwa et al, 2023). Previously, it was shown that in α-SYN protofibrils-treated astroglia, pFAK moved from the PM to the nucleus, inhibiting ROCK2 pathway that affects the actin cytoskeleton modulation and TNT formation (Raghavan *et al*, 2024). This study observes a reduction in the nuclear flatness index, indicating decreased actin stress and TNT formation after ROCK inhibitor (y-27632) treatment. In a similar vein, our RNA sequencing shows that several key regulators of FAK/Rho/ROCK are downregulated in senescent non-adherent cells at early time points (3 and 6 hours) when TNT forms. Heat maps of pathway enrichment data from RNA sequencing also detect upregulation of DNA repair and integrin-mediated cell adhesion genes in 3-hour α-SYN protofibrils-treated senescent cells, indicating cellular defence mechanisms that protect astroglia from proteotoxicity.

The actin cytoskeleton is a crucial component of TNT structure. TNTs are essential in various cellular functions, including promoting cancer cell growth, preventing apoptosis, supporting cell survival, and transmitting mechanical signals (Raghavan *et al*, 2021). Using STRING analysis combined with RNA data, we identified interacting proteins showing strong correlations among genes linked to p21/p53-driven senescence, DNA repair, the FAK/Rho/ROCK pathway, and integrin-mediated adhesion. KEGG pathway analysis highlights their crucial roles in the Hippo signaling pathway, as well as in AGE-RAGE signaling, PI3K-AKT, cytokine-receptor interactions, and Rap1 signaling pathways. The significance of actin-based machinery lies in its ability to generate both pushing and pulling forces, which are instrumental in maintaining mechanical homeostasis during biochemical changes. Cells with reduced tension activate the Hippo kinase cascade, in which YAP/TAZ translocates to the cytosol. While high tension suppresses it, allowing YAP/TAZ to enter the nucleus (Rausch & Hansen, 2020). Our data indicate that Hippo signaling is activated in α-SYN protofibrils-treated senescent cells with low actin tension, due to FAK/Rho/ROCK inhibition, thereby promoting TNT formation. Results reveal localization of YAP within the TNTs, together with actin polymers, which may facilitate the transduction of mechanical forces along these TNTs (Rausch & Hansen, 2020). During TNT formation, actin polymerization can restore actin tension through a feedback loop that involves shuttling inactive YAP to the nucleus. Interestingly, we observe the formation of TNTs between cells with differential actin tension: one with cytosolic YAP and decreased actin tension, and another with higher actin tension, in which YAP remains in the nucleus. This suggests that the actin tension gradient may influence both the fate and directionality of TNT formation between cells. Tension in the actin cytoskeleton governs mechanical homeostasis through Hippo signaling feedback loop, thereby preserving nuclear integrity and function, as well as influencing heterochromatin gene expression related to cell adhesion, differentiation, and proliferation (Fu *et al*, 2022).

Several researchers have studied the mechanism of TNTs formation, but they haven’t yet shown a direct link to Hippo signaling. However, both processes are regulated by mechanical signals: the Hippo pathway detects physical stress, while TNTs form in response to cellular stress and reduced actin tension. A study reveals that RASSF1A functions as a tumor suppressor by activating the Hippo pathway as a scaffold, promoting YAP1 phosphorylation and subsequent inactivation, while also inhibiting TNT formation by regulating GEFH1/Rab11, thereby maintaining cellular homeostasis (Dubois *et al*, 2018). Nucleolin regulates 14-3-3ζ mRNA and promotes cofilin phosphorylation, leading to TNT formation. The upregulation of the 14-3-3ζ protein (gene YWHAZ) modulates Hippo signaling to an active state (Dagar *et al*, 2021). TNTs are known to be regulated by other signaling cascades, such as the Wnt and MAPK pathways, which influence the Hippo pathway (Vargas *et al*, 2019; Cole *et al*, 2021). Other signaling pathways, such as AGE-RAGE signaling and PI3K-AKT—which are interconnected with Hippo signaling in our STRING analysis—have also been associated with the TNT biogenesis pathway (Lin *et al*, 2024; Cole *et al*, 2021). Amyloid-β aggregates promote TNT formation in neuronal cells via pPAK-mediated actin modulation (Dilna *et al*, 2021). pPAK acts as an essential downstream signaling molecule linking integrin-dependent cell adhesion to the regulation of the Hippo signaling pathway and TNT formation (Sun *et al*, 2012). Thrombospondin 1 modulates cell-matrix through integrin-mediated adhesion, interacts with Hippo signaling, and reported to link with the formation of TNTs (Mahadik & Patwardhan, 2023).

FAK in FAs links mechanotransduction to the actin cytoskeleton by regulating Rho/ROCK signaling and integrin-mediated cell adhesion, which are essential for mechanical homeostasis connecting actomyosin and nucleoskeleton structures, with Hippo signaling playing a key role (Nardone *et al*, 2017). Several studies suggest that ROCK-mediated modulation of actin plays a crucial role in TNT formation; however, the mechanisms underlying this process remain unclear. ROCK2 inhibition in microglia and astroglia promotes TNT-biogenesis (Raghavan *et al*, 2024; Scheiblich et al, 2021). In contrast, activation of the WNT5A-RhoA-ROCK1 pathway promotes TNT formation in adipose-derived stem cells (Chen *et al*, 2025). However, it is not yet clear how YAP shuttles and modulates Hippo pathway via ROCK-mediated actin-remodulating feedback loop to maintain mechanical stress by promoting TNTs.

In conclusion, α-SYN protofibrils-treated senescent astroglia disrupt nuclear structure, leading to cell de-adhesion and reduced actin tension, which in turn promotes TNT formation via ROCK2 inhibition pathway (Raghavan *et al*, 2024). RNA-sequence data indicate that senescent cells increase the expression of Hippo pathway proteins and components involved in integrin-mediated cell adhesion, which help sustain mechanical homeostasis and nucleon-cytoskeleton tension. Conversely, actin polymerization during TNT formation might trigger a mechanical signal, leading to increased cell adhesion through integrins to the extracellular matrix. This could suppress the Hippo signaling pathway by decreasing LATS1/2 kinase activity and promoting the nuclear localization and activity of the transcriptional co-activator YAP, thereby facilitating cellular recovery (Rausch & Hansen, 2020). This study shows that TNT formation in α-SYN protofibrils-treated senescent cells creates a feedback loop that aids the shuttling of YAP between the cytosol and the nucleus, modulating the Hippo pathway and stabilizing mechanical forces. However, the exact mechanism by which mechanical signals initiate TNT formation remains unclear. Nonetheless, the study reveals the complex interplay between nuclear structural integrity, mediated by Lamin A/C, and actin-cytoskeleton networks in maintaining mechanosensitive stress in α-SYN protofibrils-induced senescent astroglial cells. The process facilitates the transient formation of TNTs via the Hippo pathway and aids in the recovery of astrocytoma cells and primary astrocytes from α-SYN protofibrils-induced neurodegenerative stress by adapting biochemical changes.

## Authors’ contributions statement

SN and SC conceptualized the work; SC, AR (IIT, Goa), ASBK, AR (MAHE), and NS performed the experiments; AS and SB performed actin tension measurements; SC, SB, and SN designed the work; SP provided purified α-SYN protein and methodology support. SC, AR (IIT, Goa), SB, and SN validated and investigated all the formal analyses and interpretation of data; SC and SN wrote the first draft, and all authors edited and commented on the manuscript. All authors read and approved the final manuscript.

## Acknowledgements

The authors appreciate Genotypic Technology for providing support with next-generation sequencing and data analysis. We also thank Ms. B. Suma for her assistance with confocal microscopy at the JNCASR confocal facility and the MIRM-MAHE Bangalore microscope facility. We thank Mr Tapas Sahu for reading the manuscript and providing feedback. Additionally, we acknowledge the use of Grammarly software for English language editing. The authors confirm that no AI-generated work was used in preparing this manuscript.

## Ethics approval

All experimental protocols received approval and were performed following national animal use policies, regulated by the KMC, Manipal Academy of Higher Education, Manipal, India. The approved project title is “Tunneling nanotubes in rescuing neurodegenerative toxic burdens by facilitating glial cross-talk through cell-to-cell transfer and its role in reversal of senescence,” with approval number IAEC/KMC/75/2023, granted on October 13, 2023. Mice were euthanized with CO₂ asphyxiation followed by cervical dislocation to confirm death, in line with animal care guidelines.

## Consent for publication

All authors read the final version of the manuscript and approved it.

## Data and code availability

The raw RNA-seq data have been deposited at GEO. Additional data relevant to this study can be requested from the lead contact. No original code is provided in this article.

