## Supplementary Figures and Figure Legends for "Mechanical Signaling Drives Tunneling Nanotubes to Preserve Cytoskeleton Tension and Lamin Integrity Against α-Synuclein-Induced Senescence in Astroglia"

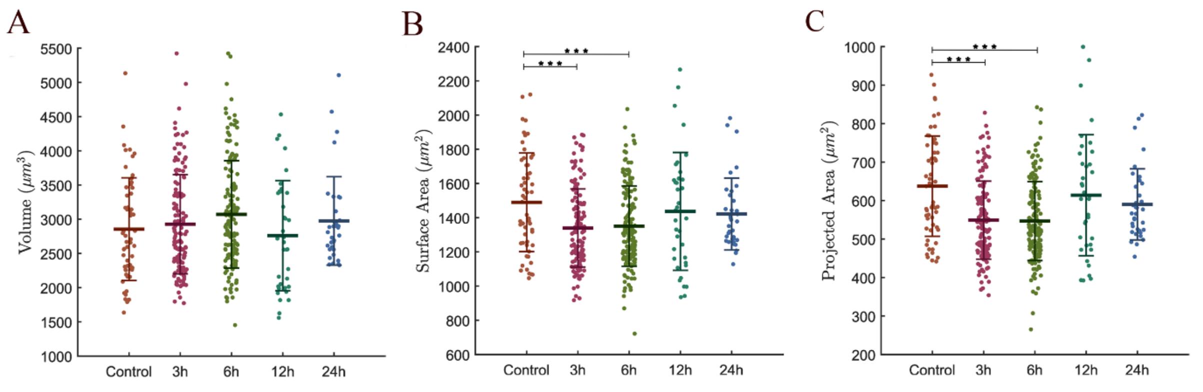


***Figure S1: Measurement of nuclear parameters.*** *Nucleus flattening (τ) and isometric scaling ($\lambda_0$) were designed to determine its shape using geometrical characteristics such as area, volume, height, and eccentricity. The analysis was performed from the Z-projection of DAPI-stained nuclei in U87MG cells. The graphs represent the quantification of A)volume, B) surface area, and C) projected area from the nucleus of U87MG cells upon α-SYN treatment (1µM) for 3, 6, 12, and 24 hours. Scale bars are denoted on the images. The graphs' data points represent the number of cells used for analysis. Data expressed as mean ± SD, ***$p \leq 0.001$. Statistics were analyzed using a two-way ANOVA. N=3.*

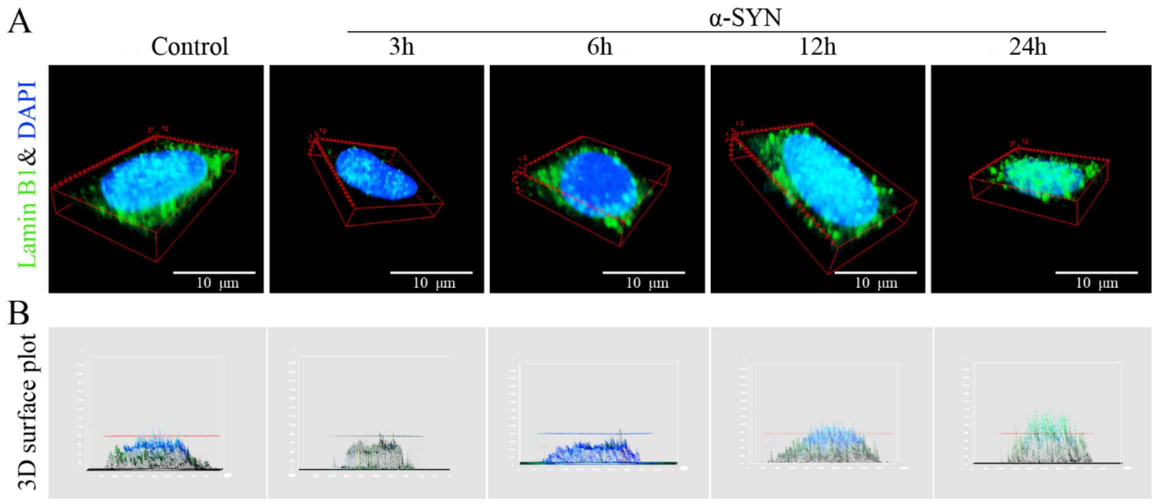


***Figure S2: Nuclear distribution of Lamin B1 upon α-SYN treatments.*** *A) 3D-view images of maximum intensity projected confocal z-stacks in U87MG cells stained with Lamin B1 (Green) and DAPI (Blue) upon α-SYN treatment (1µM) for 3, 6, 12, and 24 hours. Represented 3D surface plots showing Lamin B1 distributions in the nucleus of the α-SYN-treated cell.*

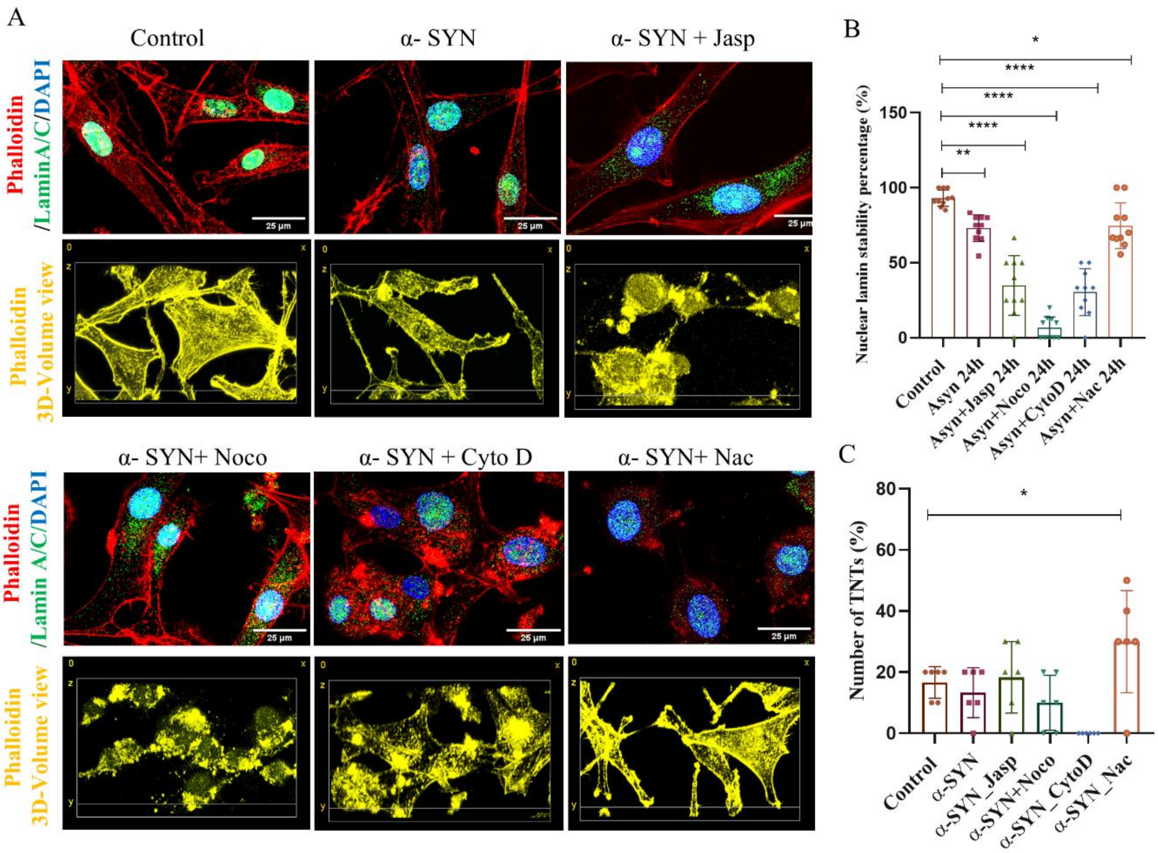


***Figure S3: Actin tension-modulating molecules inhibit TNTs and aggravate α-SYN-induced senescence.*** *A) Phalloidin (Red), Lamin A/C (Green), and DAPI (Blue)-stained U87 MG cells treated with α-SYN and α-SYN-treated cells with respective inhibitors Jasp, Noco, Cyto D and Nac at 24 hours. The small molecules inhibit TNTs, as well as disrupt the structural integrity of Lamin A/C. A) 3D-volume view stained with Phalloidin (Yellow) from superresolution microscopy shows the destruction of actin cytoskeleton networks and the inhibition of TNTs. B) Quantification of Lamin A/C intensities at the nucleus, and C) TNT numbers in percentage, after treatment with the inhibitors in the presence of α-SYN after 24 hours. The graphs' datapoints represent the number of images used for analysis, pooling data across three biological repeats. Data are expressed as mean ± SD, ***$p \leq 0.001$. Statistics were analyzed using a two-way ANOVA. N=3.*

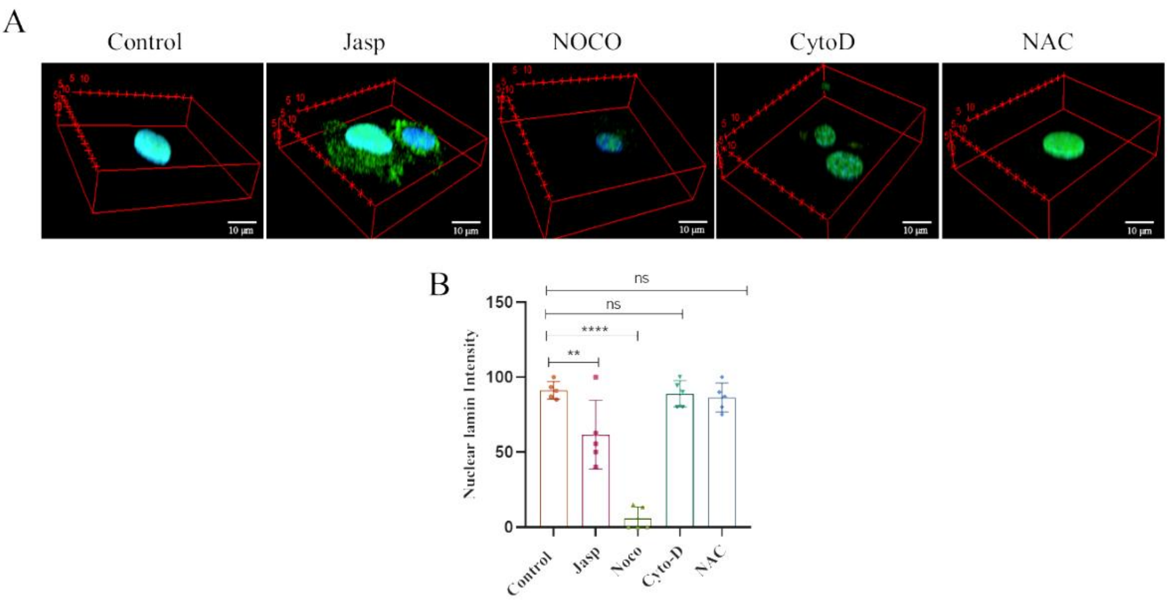


***Figure S4: Actin tension-modulating molecules in Lamin A/C distribution.*** *A) Lamin A/C (Green) and DAPI (Blue)-stained U87 MG cells treated with only small molecules Jasp, Noco, Cyto D, and Nac at 3 hours. B) Quantification of Lamin A/C intensities at the nucleus. The graphs' datapoints represent the number of images used for analysis. Data are expressed as mean ± SD,* $***p \leq 0.001$*. Statistics were analyzed using a two-way ANOVA. N = 3.*

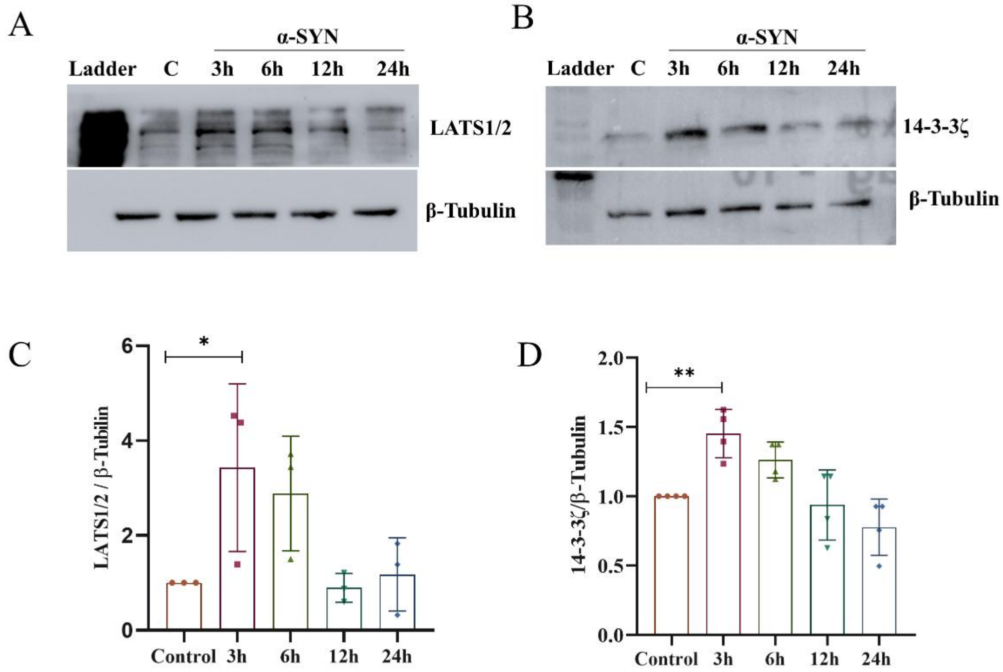


***Figure S5: LATS1/2 and 14-3-3ζ, the two key proteins in Hippo signaling, which correlate with the cytosolic translocation of YAP and senescence.*** *WB of A) LATS1/2 and B) 14-3-3ζ in*

*α-SYN-treated astroglia with time 3-24 hours. The quantification graphs show increased levels of LATS1/2 and 14-3-3ζ in α-SYN-treated astroglia at initial times 3 and 6h, Data are expressed as mean ± SD, *** $p \leq 0.001$. Statistics were analyzed using a two-way ANOVA. N = 3.*

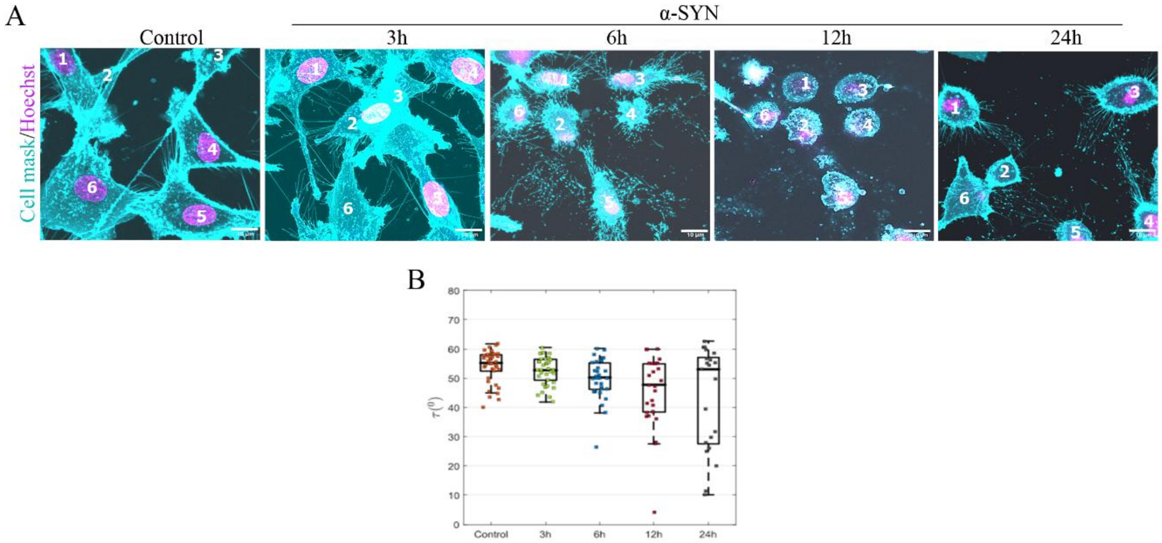


***Figure S6: Actin tension and nuclear stretch facilitating formation of TNTs.*** *A) Single cell tracking from the live cell confocal imaging using Cell Mask and (Cyan) Hoechst (Megenta)-stained U87MG cells upon α-SYN (1µM) treated cells for 3, 6, 12, and 24 hours. The images highlight significant differences in the number of TNTs at 3 hours and partial recovery of cells at 24 hours. B) Single-cell quantification of flattering index (τ) or the actin tension of U87MG cells upon α-SYN treatment (1µM) for 3, 6 ,12 and 24 hours. The graphs' data points represent the number of cells used for analysis. The result reveals that TNT-biogenesis correlates with the decrease in actin stress measured from nuclear flatness. Data are expressed as mean ± SD, *** $p \leq 0.001$. Statistics were analyzed using a two-way ANOVA. N = 3.*

**Table S1:** KEGG enrichment pathways that show significant engagement are Hippo signaling, AGE-RAGE signaling, PI3K-AKT signaling, cytokine-cytokine receptor (CCR) interaction, and the Rap1 signaling pathway

| Term description | Observed gene count | Background gene count | Strength | Signal | False discovery rate | Matching proteins in the network (labels) |
|---|---|---|---|---|---|---|
| **Hippo signaling pathway** | 7 | 154 | 1.41 | 1.58 | 3.94E-06 | *SERPINE1,AJUBA,FZD2,CCN2,GDF5,LATS2,YWHAZ* |
| **AGE-RAGE signaling pathway in diabetic complications** | 5 | 96 | 1.47 | 1.24 | 0.00015 | *MMP2,SERPINE1,ICAM1,EDN1,IL6* |
| **PI3K-Akt signaling pathway** | 7 | 349 | 1.05 | 0.92 | 0.00029 | *NGFR,THBS1,FGF18,ITGA8,YWHAZ,IL6,LPAR2* |
| **Cytokine-cytokine receptor interaction** | 5 | 282 | 1 | 0.61 | 0.008 | *NGFR,LIF,IL11,GDF5,IL6* |
| **Rap1 signaling pathway** | 4 | 201 | 1.05 | 0.54 | 0.0193 | *NGFR,THBS1,FGF18,LPAR2* |

***Figure legends of Movies:***

***Movie S1:*** *3D-volume view of Lamin A/C (red) and DAPI (blue)-stained U87 MG control cell. The nuclear volume shows an intact Lamin A/C structure encapsulating the flatter healthy nucleus.*

***Movie S2:*** *3D-volume view of Lamin A/C (red) and DAPI (blue)-stained U87 MG cell treated with α-SYN for 3 hours. The nuclear volume shows a Lamin A/C-devoid rounded nuclear structure.*

***Movie S3:*** *3D-volume view of Lamin A/C (red) and DAPI (blue)-stained U87 MG cell treated with α-SYN for 24 hours. The nuclear volume shows an intact Lamin A/C structure encapsulating a relatively flatter and healthier nucleus (Movie S1).*

***Movie S4:*** *3D-volume view of U87 MG cell stained using Cell mask (Green) treated with α-SYN for 3 hours. The image shows TNTs forming between less-adhered cells and those with relatively better adhesion.*

***Movie S5:*** *3D-volume view of primary astrocytes treated with α-SYN for 3 hours, immuno-stained with YAP (Green) and phalloidin (Red). The image depicts TNT formation between cells: one with cytosolic YAP and decreased actin tension, and another with higher actin tension, where YAP remains in the nucleus.*
